# Allele-Specific QTL Fine-Mapping with PLASMA

**DOI:** 10.1101/650242

**Authors:** Austin T. Wang, Anamay Shetty, Edward O’Connor, Connor Bell, Mark M. Pomerantz, Matthew L. Freedman, Alexander Gusev

## Abstract

Although quantitative trait locus (QTL) associations have been identified for many molecular traits such as gene expression, it remains challenging to distinguish the causal nucleotide from nearby variants. In addition to traditional QTLs by association, allele-specific (AS) QTLs are a powerful measure of cis-regulation that are largely concordant with traditional QTLs, and can be less susceptible to technical/environmental noise. However, existing asQTL analysis methods do not produce probabilities of causality for each marker, and do not take into account correlations among markers at a locus in linkage disequilibrium (LD). We introduce PLASMA (PopuLation Allele-Specific MApping), a novel, LD-aware method that integrates QTL and asQTL information to fine-map causal regulatory variants while drawing power from both the number of individuals and the number of allelic reads per individual. We demonstrate through simulations that PLASMA successfully detects causal variants over a wide range of genetic architectures. We apply PLASMA to RNA-Seq data from 524 kidney tumor samples and show that over 17 percent of loci can be fine-mapped to within 5 causal variants, compared less than 2 percent of loci using existing QTL-based fine-mapping. PLASMA furthermore achieves a greater power at 50 samples than conventional QTL fine-mapping does at over 500 samples. Overall, PLASMA achieves a 6.9-fold reduction in median 95% credible set size compared to existing QTL-based fine-mapping. We additionally apply PLASMA to H3K27AC ChIP-Seq from 28 prostate tumor/normal samples and demonstrate that PLASMA is able to prioritize markers even at small samples, with PLASMA achieving a 1.3-fold reduction in median 95% credible set sizes over existing QTL-based fine-mapping. Variants in the PLASMA credible sets for RNA-Seq and ChIP-Seq were enriched for open chromatin and chromatin looping (respectively) at a comparable or greater degree than credible variants from existing methods, while containing far fewer markers. Our results demonstrate how integrating AS activity can substantially improve the detection of causal variants from existing molecular data and at low sample size.

## 1 Introduction

A major open problem in genetics is understanding the biological mechanisms underlying complex traits, which are largely driven by non-coding variants. A widely adopted approach for elucidating these regulatory patterns is the identification of disease variants that also modify molecular phenotypes (such as gene expression) [1–4]. These variants, known as quantitative trait loci (QTLs), are typically single nucleotide polymorphisms (SNPs) that exhibit a statistical association with overall gene expression abundance [5–8]. Although QTL association analysis is now mature, it remains challenging to identify the precise variants that causally influence the molecular trait (as opposed to variants in linkage disequilibrium (LD) with causal variants), a task known as fine-mapping [9]. As only a small subset of QTL-associated markers are estimated to be causal [10, 11], direct experimental validation is prohibitive and has motivated statistical fine-mapping solutions [12]. The aim of statistical fine-mapping is to quantify the probability of each marker being causal, allowing one to prioritize the most likely causal markers, and thus formally quantify the effort needed for experimental validation. Recent statistical fine-mapping methods operate on summary QTL statistics and can handle multiple causal variants by modeling the local LD structure [13–16]. These models have two outputs to help guide the prioritization of putative causal SNPs. First, for each marker, a Posterior Inclusion Probability (PIP) is calculated, which are corresponds to the marginal probability of causality for the given marker. Second, a *n*%-confidence credible set is created: a set of markers with an *n*% probability of containing all the causal markers. Although QTL studies have sufficient power to identify thousands of associations, they are typically insufficient for fine-mapping below dozens of credible variants, even for very large studies [5, 17]. The need for large studies is severely limits QTL analyses of expensive assays such as ChIP or single-cell RNA-seq, or of difficult to collect tissues.

Here, we sought to improve molecular fine-mapping by leveraging intra-individual allele-specific (AS) signal, which is a measure of cis-regulatory activity that is independent of total, interindividual variation. For heterozygous variants residing in expressed exons, it is often possible to map expressed reads to each allele and quantify the extent that molecular activity is allelespecific [6, 18–21]. AS analysis allows for a precise comparison of the effects on molecular activity that are specific to each allele (cis-effects), while controlling for effects affecting both alleles (trans-effects). Thus, AS data is inherently less noisy than regular QTL data, which captures total phenotype regardless of source. Allele-specific data has furthermore been used to quantify cis-regulation, implying that both AS and regular QTL features represent the same underlying cis-regulatory patterns [22]. Several methods have recently been developed to robustly identify asQTLs [19, 20, 23], but the calculated association statistics follow a different distribution than QTL summary statistics and cannot be directly integrated into existing fine-mapping software to produce valid posterior measures and credible sets.

To combine the scalability of QTL analysis with the power of AS analysis, we introduce PLASMA (PopuLation Allele-Specific Mapping), a novel fine-mapping method that gains power from both the number of individuals and the number of allelic reads per individual. By modeling each locus across individuals in an allele-specific and LD-aware manner, PLASMA achieves a substantial improvement over existing fine-mapping methods with the same data. We demonstrate through simulations that PLASMA successfully detects causal variants over a wide range of genetic architectures. We apply PLASMA to diverse RNA-Seq data and ChIP-seq data and show a significant improvement in power over conventional QTL-based fine-mapping.

## 2 Methods

### 2.1 Overview of PLASMA

PLASMA’s inputs are determined from a given individual-level sequencing-based molecular phenotype (gene or peak) and the corresponding local genotype SNP data (Figure 1a). For each sample, we assume the variant data is phased into haplotypes and expression reads have been mapped to each variant. Reads intersecting heterozygous markers (signified as fSNPs, or feature SNPs, indicated with green or purple on the figure) are then assigned to a particular haplotype, indicated as blue or red on the figure. These reads are then aggregated in an haplotype-specific manner to produce a total expression phenotype and an allelic imbalance phenotype. This aggregation of reads is analogous to the way existing methods such as RASQUAL and WASP calculate allelic fractions and total fragment counts [19, 20]. The total expression phenotype (*y*) is simply the total number of mapped reads. The allelic imbalance phenotype (*w*) is defined as the log read ratio between the haplotypes. This log-odds-like phenotype has previously been used to analyze asQTL effect sizes, showing consistency with conventional QTL analysis [22]. To mitigate the effect of mapping bias, we run state of the art mapping bias and QC pipelines on all RNA-Seq and ChIP-Seq data prior to analysis [19].

**Figure 1:**
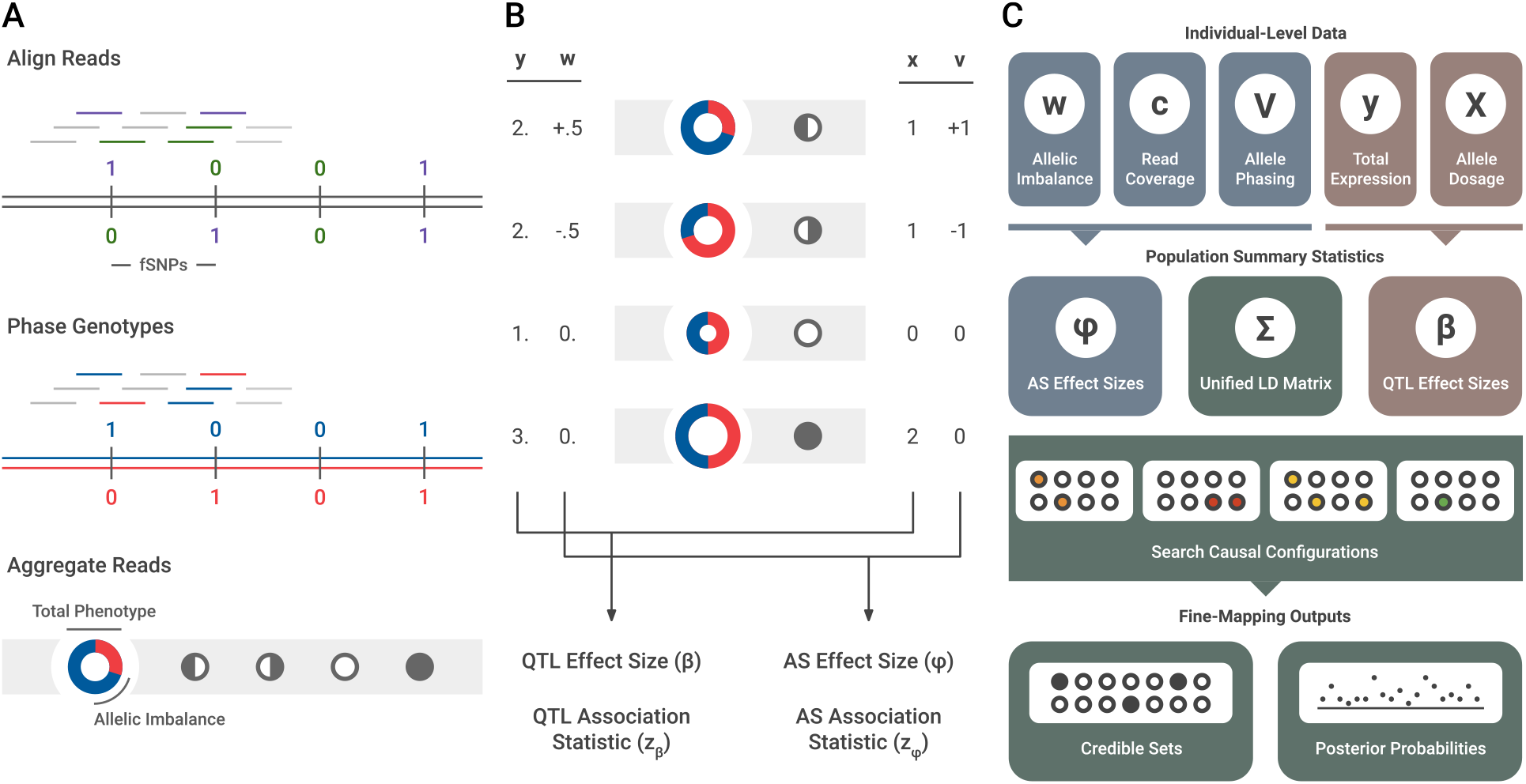
Overview of the PLASMA method. (a) Pre-processing of sequence-based data. First, reads are mapped to the sample’s genotype. Reads intersecting markers are colored. Then, the sample’s genotype is phased. Reads intersecting heterozygous markers can then be mapped to a particular haplotype. Lastly, reads across the locus are aggregated in an allele-specific manner. To visualize this data, the expression is represented by a ring chart, and the genotypes by pedigree symbols. In the ring chart, the diameter signifies the total read count, and the colors signify the proportion of reads coming from each haplotype. For the pedigree symbols, a white circle signifies a wild-type homozygote, a shaded circle signifies a alternative homozygote, and a half-shaded circle signifies a heterozygote. In heterozygotes, the direction of shading corresponds to the direction of heterozygosity (phasing). (b) Visual representation of QTL and AS statistics under a single causal variant,where the alternative allele increases expression. The total expression (*y*) is determined by the allelic dosage (*x*), whereas the allelic imbalance (*w*) is determined by the phasing (*v*). These two sets of data are used to calculate QTL and AS association statistics (*z_β_* and *z_ϕ_*). (c) Diagram of PLASMA’s fine-mapping process. First, QTL and AS statistics are calculated from read data. Then, these statistics, along with an LD matrix, are used to generate probabilities for causal configurations. By searching through the space of these causal configurations, the model produces credible sets and posterior probabilities for each marker.

PLASMA integrates two statistics computed for each marker to perform fine-mapping: a QTL association statistic (*z_β_*) based on the total phenotype and an AS association statistic (*z_ϕ_*) based on the allelic imbalance phenotype. Figure 1b shows how a causal marker influences total expression and allelic imbalance, and how this effect influences the statistics for the marker (see Methods for quantitative explanations of the statistics). Here, the causal marker’s alternative allele causes higher expression compared to the wild-type allele. Increasing the dosage (*x*) of the alternative allele increases the total expression (*y*) at the locus. The effect size (*β*), consistent with a typical QTL analysis, quantifies the association between a marker’s allelic dosage and the total expression at the locus with a linear relationship with residuals *ϵ*:

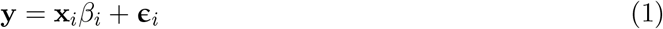

From this effect size, PLASMA calculates *z_β_*, the QTL association statistic (See Methods for the precise relationship with *β*). Note that this statistic is not dependent on haplotype-specific data.

On the other hand, looking at the heterozygotes, the haplotype possessing the alternative allele has a higher expression than the haplotype possessing the wild-type allele. In other words, the direction of imbalance of expression (*w*) is the same as the phasing (*v*) of the allele. The *ϕ* effect size quantifies the association between a marker’s phasing with the imbalance of expression. An important departure from existing methods is that PLASMA models a linear relationship between the phase of a causal marker and the log read ratio, rather than directly relating the genotype to the allelic fraction in a non-linear manner with residuals *ζ*:

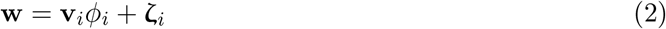

To calculate the AS association statistic *z_ϕ_*, PLASMA models the quality of each sample, taking into account each sample’s read coverage and read overdispersion (Figure 1c, see Methods for the precise weighing scheme).

These QTL and AS association statistics, together with the local LD matrix, are then jointly used to fine-map the locus (Figure 1c, see Methods for the fine-mapping model). Since PLASMA models both *z_β_* and *z_ϕ_* as a linear combination of genotypes, *z_β_* and *z_ϕ_* have identical LD (see Appendix for proof). PLASMA assumes that the QTL and AS statistics measure the same underlying cis-regulatory signal and are thus expected to have the same direction and same causal variants (but see Discussion for possible model violations). Although they both measure regulatory effects, the two statistics have independent noise because the haplotype-level variance within individuals is considered only in AS analysis, allowing them to be used jointly in fine-mapping. Furthermore, PLASMA accepts, as a hyperparameter, a correlation between QTL and AS effects, allowing the two sets of statistics to utilize a joint probability distribution (though our analyses show that setting this hyperparameter to zero yields the most power). The distribution is used to assign a probability to a given causal configuration, a binary vector signifying the causal status of each marker in the locus. Although the correlation between QTL and AS causal effects can vary based on the hyperparameter specification, PLASMA assumes that the AS and QTL phenotypes have the same causal variants. PLASMA searches through the space of possible causal configurations, within a constraint on the number of causal variants. This procedure is related to that in CAVIAR, CAVIARBF, and FINEMAP [13–15], but generalized to the two correlated expression phenotypes. From these scored configurations PLASMA computes a posterior inclusion probability (PIP) for each marker, indicating the marginal probability that a marker is causal, and a *ρ*-level credible set containing the causal variant with *ρ* probability.

### 2.2 Modeling QTL and AS summary statistics

Marginal QTL effect sizes for a given locus are calculated under the conventional linear model of total gene expression, with the allelic dosage (**x**) as the independent variable, and the total expression (**y**) as the independent variable. Let us consider a QTL study of a given locus with *n* individuals and *m* markers. Let y be an (*n* × 1) vector of total expression across the individuals, recentered at zero. Given a marker *i*, let x_*i*_ be a zero-recentered vector of dosage genotypes. The genetic effect *β_i_* of marker *i* on total gene expression is defined as follows:

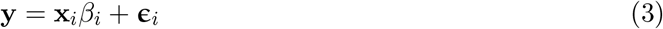

The empirical value of *β_i_* is determined with the maximum likelihood estimator, equivalent to the ordinary-least-squares linear regression estimator:

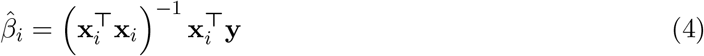

The QTL summary statistic (Wald statistic) for marker i is defined as:

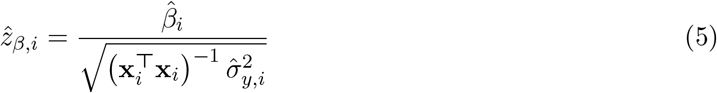

where 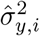 is calculated from the residuals.

AS effect sizes are calculated under a weighted linear model, with the phasing (**v**) as the independent variable, and the allelic imbalance (**w**) as the dependent variable. PLASMA models allele-specific expression under the observation that a cis-regulatory variant often has a greater influence on the gene allele of the same haplotype. A marker’s phase *υ* is 1 if haplotype *A* contains the alternative marker allele, −1 if haplotype *B* contains the alternative marker allele, and 0 if the individual is homozygous for the marker. Let *w* be the log expression ratio between haplotypes A and B, *ϕ_i_* be the AS effect size of variant *i*, and *ζ_i_* be the residual, interpreted as the log baseline expression ratio between haplotypes A and B. Additionally, a sampling error *τ_j_* = *ŵ_j_* – *w_j_* is defined for each individual, quantifying the quality of data from the sample. The genetic effect of marker i on allele-specific expression is as follows:

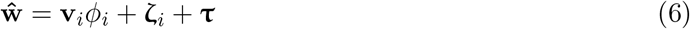

Experimentally-derived AS data, such as RNA-Seq data, yield reads that are mapped to a particular haplotype. For a given individual *j*, let *c_A,j_* be the allele-specific read count from haplotype *A*. The allele-specific read count is modeled as drawn a beta-binomial distribution, given the total mapped read count *c_j_*:

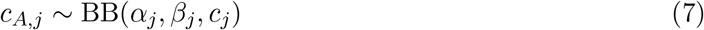

This beta binomial model is used to estimate the variance of the sampling error *τ_j_*:

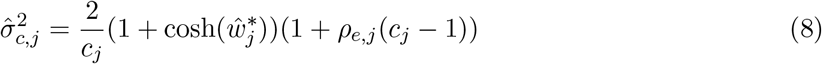

where *ρ_e,j_* is the overdispersion and 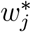 is an adjusted estimator of *W_j_* to reduce the bias of 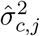. (Full derivation in Supplementary Methods).

Due to heteroscedasticity among individuals, the AS effect size *ϕ_i_* is estimated in a weighted manner, giving larger weights to individuals with lower estimated sampling error. Given individual *j*, the weight for *j* is set as the inverse of the estimated sampling error variance:

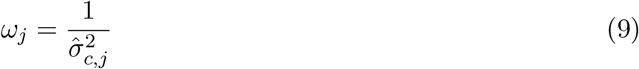

Let weight matrix **Ω** be a diagonal matrix with **Ω**_*j,j*_ = *ω_j_*. We use the weighted-least-squares estimator for *ϕ_i_*:

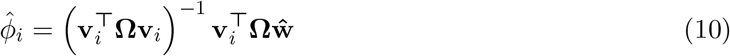

With this estimator, the AS association statistic for marker *i* is calculated as the AS effect size divided by the estimated variance of the effect size (full derivation in Appendix):

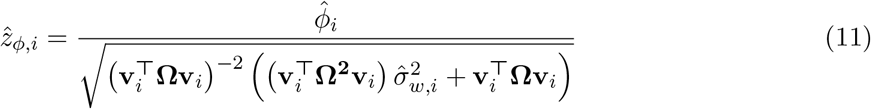

In the case of computationally phased data, there may exist phasing errors that would decrease the accuracy of the estimated effect sizes 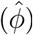. With imperfect phasing, the observed phasing 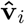 may differ from the true phasing **v**_*i*_, a modified equation may be used to calculate the AS z-score given the per-SNP probability of mis-phasing *ψ_i_*:

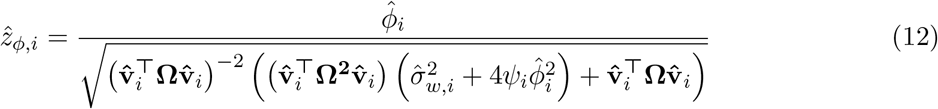

Fine-mapping simulation results under imperfect phasing are presented in the Appendix and in Figure S5.

### 2.3 Inference of credible sets and posterior probabilities

PLASMA defines a joint generative model for total (QTL) and haplotype-specific (AS) effects on expression. Let 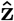 be the combined vector of AS association statistics and QTL association statistics:

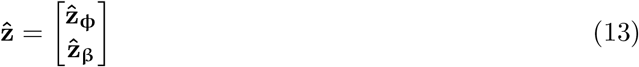

Let **R**_z_ be the genotype LD matrix, and *r_βϕ_* be a hyperparameter describing the overall correlation between the QTL and AS summary statistics calculated across all loci. Let the combined correlation matrix **R** as:

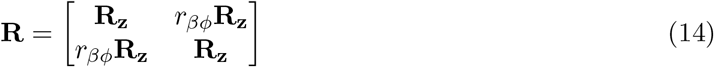

The joint z-scores are modeled under a multivariate normal distribution, with covariance **R**:

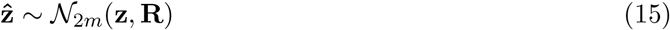

PLASMA utilizes a likelihood function that gives the probability of statistics 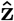, given a causal configuration. Let a causal configuration **c** be a vector of causal statuses corresponding to each marker, with 1 being causal and 0 being non-causal. PLASMA assume that the causal configuration is the same for the QTL and AS signals.

Let hyperparameters 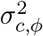 and 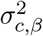 be the variance of AS and QTL causal effect sizes, respectively. Furthermore, let the jointness parameter *r_c,βϕ_* be the underlying correlation of the causal QTL and AS effect sizes. (This is not to be confused with *r_βϕ_*, which concerns the correlation between the observed association statistics. See the Appendix for a mathematical relationship between these two hyperparameters.) These three hyperparameters are closely related to the heritability of gene expression (see Appendix). Let Σ_c_ be the covariance matrix of causal effect sizes given a causal configuration:

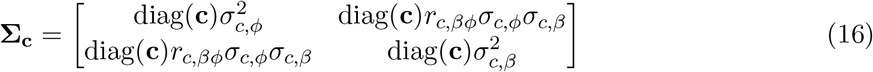

PLASMA’s likelihood for a causal configuration is defined as:

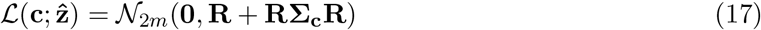

Let *γ* be the prior probability that a single variant is causal and 1 – *γ* as the probability that a variant is not causal. The prior probability of a configuration consisting of m variants is defined as:

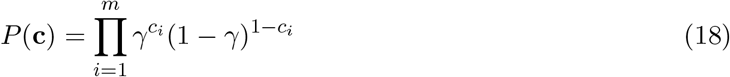

With the prior and likelihood, the posterior probability of a causal configuration, normalized across the set of all possible configurations ℂ is calculated as:

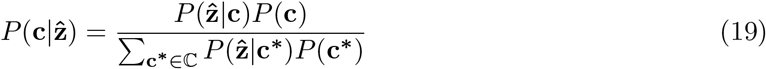

PLASMA defines the *ρ*-level credible set 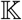 as the smallest set of markers with a *ρ_c_* probability of including all causal markers. Let 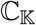 be the set of all causal configurations whose causal markers is a subset of 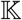, excluding the null set. The credible set confidence level *ρ_c_* is calculated as the sum of the probabilities of the configurations in 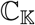:

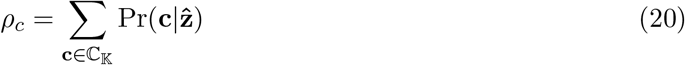

Additionally, PLASMA defines a marker’s posterior inclusion probability (PIP) as the probability that a single given marker is causal, marginalized over all other markers. This probability is calculated by summing over all configurations containing the marker.

To reduce the number of configurations to evaluate in the case of multiple causal variants, PLASMA usees the heuristic that configurations with significant probabilities tend to be similar to each other. PLASMA uses a shotgun stochastic search procedure to find all configurations with a significant probability. For each iteration of the algorithm, the next configuration is drawn randomly from the neighborhood of similar configurations, weighted by the posterior probability of each candidate. The search is terminated under the preseumption that all configurations with nonzero probability have been uncovered.

Given the large number of configurations evaluated, it is impractical to calculate the best possible credible set satisfying *ρ_c_*. Instead, PLASMA uses a greedy approximation algorithm. At each step, before *ρ_c_* is reached, the algorithm adds the marker that increases the confidence the most.

### 2.4 The Jointness Parameter in PLASMA

Although PLASMA always assumes the same causal variants for QTL and AS, the correlation between QTL and AS causal effects can be set in PLASMA-J with a jointness hyperparameter *r_c,βϕ_*. A high value (near 1) assumes that the QTL and AS causal effects tend to be consistent in magnitude, while a low value (near zero) assumes more disparity. Note that this is unrelated to choice of causal variants, and PLASMA assumes that QTL and AS share the same causal variants regardless of the jointness parameter.

A previous analysis comparing QTL effects with a similar formulation of AS effects has uncovered a highly nonlinear relationship, especially with QTL effects calculated using untransformed total expression data [22]. As a futher complication, this relationship between QTL and AS effects is shown to be highly dependent on allele frequency. Thus, even under the assumption that QTL and AS signals share a causal variant, there is no guarantee of a strong linear correlation between QTL and AS effect sizes. Due to this uncertaintly, the jointness parameter to zero by default, making no assumption on the relationship between QTL and AS effect sizes.

To empirically evaluate the effect of the jointness parameter on fine-mapping performance, PLASMA-J was run with different values of the jointness parameter on simulated loci. Figure S2 shows the distribution of PLASMA-J credible sets with different values of jointness, ranging from 0 to 0.99. In the one causal variant case, results are largely invariant to the parameter below a value of 0.99.

### 2.5 Generation of simulated loci

Genotype data was sampled from phased SNP data using the CEU population in the 1000 Genomes Project. First, a contiguous section of markers is randomly chosen. Next, a random selection of samples are randomly selected from the section. The genotypes corresponding to the chosen samples yield two haplotype matrices, denoted as **H**_*a*_ and **H**_*b*_.

Among the markers, the desired number of causal markers is randomly selected. In the case of multiple causal variants, each causal marker is assigned a relative effect size, sampled from a normal distribution with zero mean and unit variance. For each individual, the ideal un-scaled gene expression for each haplotype **q**_*a*_ and **q**_*b*_ is determined by multiplying the relative effect sizes with each haplotype matrix.

Read count data is simulated with this haplotype-specific expression. In real data, only a fraction of the reads can be mapped to a specific haplotype. Due to this difference between total reads and mapped reads, the allelic imbalance and the total read count (QTL) are calculated separately.

To calculate total read count data, the total ideal un-scaled expression **q**_*t*_ is defined as **q**_*a*_ + **q**_*b*_, the sum of the haplotype-specific un-scaled gene expression. Gaussian-distributed noise is then added so that the variance of **q**_*t*_ is consistent with the total variance across samples as specified by the QTL heritability. Finally, this final expression is scaled so that the total expression across samples is of unit variance. Total read counts are not explicitly generated, since a multiplicative factor across samples does not influence the QTL association statistics calculated by the model. This is reflective of typical QTL study protocols which aggressively rank/quantile normalize the data to fit a normal distribution.

To calculate allele-specific read counts, heritability, mean read coverage, and the total variance of the AS phenotype are taken into account. The ideal allelic imbalance phenotype is determined as logit 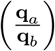 (calculated element-wise). Gaussian-distributed noise is then added so that the signal-to-noise ratio of the phenotype’s variance is consistent with the specified AS heritability. This noisy phenotype is then scaled to the specified total variance. The read coverage for each sample is then drawn from a Poisson distribution, given the mean read coverage. Lastly, allele-specific read counts are generated from these phenotypes, with the counts for each sample being drawn from a beta-binomial distribution.

### 2.6 Comparison of existing models with PLASMA

Our analyses benchmark PLASMA against existing fine-mapping methods. Two distinct versions of PLASMA are tested, “PLASMA-J” and “PLASMA-AS.” The PLASMA-J (Joint-Independent) version looks at both AS and QTL statistics, assuming a shared set of AS and QTL causal variants, and also that the AS and QTL causal effects are uncorrelated. The “PLASMA-AS” version is restricted to only AS data. As a baseline, we compare PLASMA to a QTL-Only version of PLASMA and to the CAVIAR method (expected to be equivalent to PLASMA QTL-Only) [13]. The behavior and performance of CAVIAR is representative of similar QTL-based methods such as CAVIARBF, FINEMAP, and PAINTOR without functional annotation data [14–16]. The versions of PLASMA are furthemore compared against the only other publicly-released fine-mapping method (to our knowledge) that integrates AS data described in the pre-print of Zou et al., 2018 [24]. This unnamed method, denoted as “AS-Meta,” utilizes the association between SNP heterozygosity and a binary indicator of allelic imbalance. By binarizing allelic imbalance, AS-Meta is expected to lose power relative to treating imbalance as a quantitative phenotype but may be more robust to spurious AS signal. Furthermore, AS-Meta utilizes only indicators of heterozygosity, rather than marker phasing. AS-Meta can therefore be used with unphased genotypes, but at the expense of being unable to leverage the direction of the allelic effect. Lastly, as an additional comparison with an AS-based method, we analyze the performance of RASQUAL, a method for inferring alllele-specific genetic effects using both allelic and total expression signal. Note that RASQUAL only computes allele-specific effect sizes for each marker and is not intended to compute credible sets or posterior marginal probabilities. Traditional fine-mapping on RASQUAL statistics is made possible by converting RASQUAL chi-square statistics back to quasi-z-scores with sign based on the direction of the RASQUAL effect-size. These statistics are then fed into standard QTL-only fine-mapping to obtain credible sets and posterior probabilities. We denote the modification of RASQUAL as “RASQUAL+”. This process is comparable to fine-mapping using a combined AS/QTL effect, rather than modeling QTL and AS effects separately.

### 2.7 Quality control of genotype data

For TCGA data, germline genotype calls are downloaded from the Genomic Data Commons. For PrCa ChIP samples, germline genotypes are called from blood (personal communication). Genotypes were imputed to the Haplotype Reference Consortium [25] using the Michigan Imputation Server [26] and restricted to variants with INFO greater than 0.9 and MAF greater than 0.01. Variants are further restricted to QC-passing SNPs from Ref. [25] which represent common, well-mapped variants from the 1000 Genomes project.

### 2.8 Quality control of RNA-seq data

Raw RNA-seq BAM files are downloaded from the Genomic Data Commons. Initial RNA-seq mapping and alignment are performed following TCGA parameters for the STAR aligner [27]. Mapping bias is accounted for by re-mapping using the WASP pipeline [19] and the STAR aligner with the same parameters. Reads are randomly de-duplicated as recommended by the WASP pipeline.

Somatic copy number calls are downloaded from FireBrowse and local beta-binomial overdispersion parameters are estimated for each contiguous region of copy number change.

### 2.9 Quality control of ChIP-seq data

ChIP-seq reads are aligned using bwa and default parameters [28], and peaks are called using MACS2 and default parameters (with DNA-seq input provided as control) [29]. Peaks are then unified across all samples. Mapping bias is accounted for by re-mapping using the WASP pipeline and the bwa aligner with the same parameters. Reads are randomly de-duplicated as recommended by the WASP pipeline. Beta binomial overdispersion parameters are estimated globally for each sample as somatic copy number was expected to be minimal.

### 2.10 Allele-specific quantification

The StratAS algorithm is used to quantify allele-specific signal and identify initially significant features for fine-mapping [23]. For each peak/gene (the feature) and individual all reads at heterozygous SNPs in the feature are aggregated to compute the haplotype-specific read counts, and summed across the two haplotypes of each individual to compute the QTL read counts. Each QC passing variant within 100kb of the feature are then tested for an allele-specific association with the feature and features significant at a genome-wide false discovery rate (FDR) of 5% are retained for fine-mapping.

### 2.11 Functional enrichment analysis

For QTLs fine-mapped from RNA-seq we select regions of accessible chromatin in the most relevant tissue as reference the functional feature, reasoning that high-confidence causal variants should be more abundant in accessible regions. For QTLs fine-mapped from ChIP-seq we select chromosome looping anchors from Hi-ChIP in the relevant tissue as the reference functional feature, reasoning that high-confidence causal variants should be more abundant in regions that are in conformation with promoters.

Enrichment is then estimated by computing the proportion of markers in credible sets that intersect with the functional feature. Controls are calculated as the intersection between all tested markers and the functional feature. Odds ratios and p-values are computed with Fisher’s exact test.

## 3 Results

### 3.1 Simulation framework

We evalulate PLASMA with a framework that simulates the expression of whole loci in an allelespecific manner. This simulation framework jointly simulates total reads and allele-specific read counts, under given values of the number of causal variants, the QTL heritability, the AS heritability, the variance of the AS phenotype across samples, and the expected read coverage (see Methods). The variance and heritability of the AS phenotype are handled by two separate parameters, where the former describes the total spread of allelic imbalance, and the latter specifies the fraction of the variance that is due to genetic effects. This allows us to investigate cases where a significant amount of observed imbalance is caused by non-genetic variance in the allelic expression. To quantify the total variance of the AS phenotype in the population, we define the “standard allelic deviation” (d) as the standard deviation of the AS phenotype w, quantified on the allelic fraction scale (between 0.5 and 1). Importantly, this quantity is independent of the genetic effect, which is controlled by the heritability parameter. Simulations were performed using real phased haplotype data from the 1000 Genomes Project European samples. Parameter settings for simulation analyses are shown in Table S1.

As the performance of standard QTL association models is well established, we first focused on performance of our proposed AS statistic. Figure S3a shows how the mean *z_ϕ_* varies as a function of standard allelic deviation and mean read coverage at a fixed AS heritability of 0.5. Second, Figure S3b shows how the mean *z_ϕ_* varies as a function of standard allelic deviation and heritability with mean coverage fixed at 100. The statistic is the greatest at high read coverage and high heritability, consistent with the degree of experimental and intrinsic signal available to the model. These results hold even at low AS variance (*d* = 0.6) and show that PLASMA does not conflate high AS variance (standard allelic deviation) with high signal (coverage or heritability). This robustness to variance in the AS phenotype makes the model resistant to false-positives driven by non-genetic sources of allelic variance. At very high variance (*d* > 0.8), *z_ϕ_* shows a sharp decrease. This decrease in signal is due to an increase in the sampling error of the AS phenotype (*w*) at high overall variance, as shown in Equation 47 (See Appendix for a mathematical relationship between total variance and sampling error.)

### 3.2 Comparison with existing methods in simulation

First, we evaluate how well each PLASMA prioritizes candidate causal markers using simulated loci with one causal variant, compared to existing QTL and AS-based methods. We define the “inclusion curve” for each model, where markers are ranked by posterior probability and added one by one to a cumulative set (note that this set is not dependent on the definition of a credible set). The x axis represents the cumulative number of markers chosen, and the y axis represents the “inclusion rate,” the proportion of true causal markers among the chosen markers. Figures 2a and d show inclusion plots at low and high AS variance, respectively. As expected, FINEMAP, QTL-Only, and the CAVIAR methods are indistinguishable and do not vary with AS variance. Due to this similarity in results, FINEMAP is used as the primary QTL-based methods in subsequent analyses. Furthermore, PLASMA-J and PLASMA-AS perform similarly at both levels of AS variance. Additionally, AS-Meta’s performance exhibits a dependency on the degree of AS variance. Lastly, RASQUAL+ at high AS variance does significantly improve over QTL-based methods, but not as well as PLASMA. At low AS variance (with same amount of signal and noise), RASQUAL+ performs considerably worse, indicating that RASQUAL+ is more sensitive to the genetic architecture of the locus than PLASMA is.

**Figure 2:**
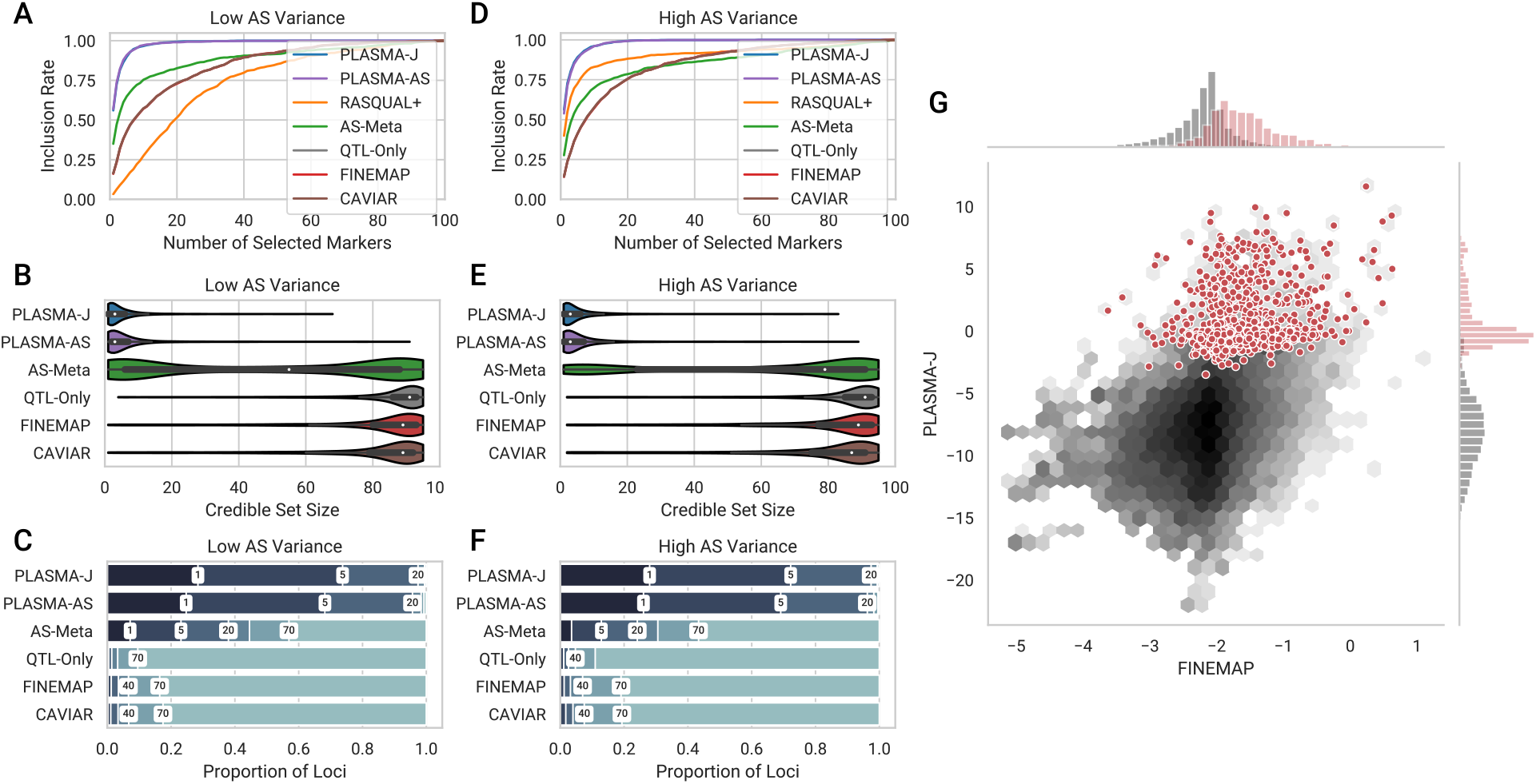
Comparison of fine-mapping methods on two sets of simulated loci of 100 markers and 100 samples each. These two sets differ in the variance of the AS phenotype, but with a fixed mean coverage and AS heritability. (a, d) Inclusion curve at low and high AS variance, respectively (0.6, 0.8 standard allelic deviation). The y-axis shows the expected proportion of markers included in the credible set, and the x-axis shows the number of selected markers by posterior probability. (b, e) Distribution of 95% confidence credible set sizes at low and high AS variance, respectively. (c, f) Proportions of loci with 95% credible set sizes within given thresholds at low and high AS variance, respectively. Thresholds used are 1, 5, 20, 40, 70, and 100 markers. (g) A per-snp fine-mapping comparison between PLASMA-J and FINEMAP across 500 simulated loci with one causal variant each. The axes denote the posterior log-odds of causality for FINEMAP and PLASMA-J, respectively. The black hexagons represent the joint distribution of all markers, while the red dots represent specifically the causal markers. Univariate histograms for PLASMA-J and FINEMAP are plotted along the margins.

Second, we evaluate the ability of each model to rule out likely non-causal markers in simulated loci with one causal variant. We conduct a direct comparison of the distributions of the 95% confidence credible sets, with smaller sets indicating higher specificity. Figures 2a and d show distribution plots at low and high AS variance, respectively. At low variance, PLASMA-J offers the smallest median credible 95% set size (3.0), followed by PLASMA-AS (3.0), then AS-Meta (55.0), and lastly the QTL-based methods: FINEMAP (89.0), CAVIAR (89.0), and QTL-Only (91.0). There is some variation due to differences in calibration among the methods, but all QTL-based methods have recall above 0.95. PLASMA appears robust to changes in AS variance; at high AS variange, medians are 3.0 for PLASMA-J and 3.0 for PLASMA-AS. In contrast, the performance of AS-Meta varies significantly with the degree of AS variance, even when the underlying signal (coverage and heritability) is constant, with a median set size of 79.0 at high variance. This sensitivity may be due to the fact that AS-Meta does not incorporate marker phasing, and thus must rely solely on the intensity of imbalance, rather than the direction of imbalance. Here, RASQUAL+ does not generate meaningful credible sets, with 95% credible set recall being 0.06 and 0.58 for low and high AS variance, respectively. RASQUAL+ is therefore not included in further fine-mapping analyses, though we underscore that RASQUAL remains an effective tool for QTL discovery.

Third, we directly compare how PLASMA-J and FINEMAP prioritize a common set of variants pooled from 500 loci, each with 100 total markers and one causal marker. Figure 2g shows a joint histogram of log posterior marginal odds of these 50000 variants, with causal variants highlighted in red. Distributions of posterior log-odds for each method are shown as univariate histograms along each axis. As expected, PLASMA and FINEMAP posterior log-odds are positively correlated. Comparing the distribution of the odds of causal variants to those of the rest, it is furthermore evident that PLASMA more confidently assigns probabilities of causality, and can much more cleanly segregate causal from non-causal variants.

Lastly, we run the AS-based methods across a wide range of coverage and heritability conditions, recording the mean 95% confidence credible sets, shown in Figure 3. Figures 3a-c show mean credible set sizes as a function of AS variance and coverage, and Figures 3d-f show mean credible set sizes as a function of AS variance and AS heritability. In terms of the range of set sizes, PLASMA-J performs the best (3.2 markers on average at best conditions), followed by the PLASMA-AS (3.4 at best conditions), and lastly the AS-Meta method (31 at best conditions). Generally speaking, all methods show results consistent with the behavior of *z_ϕ_* in Figure S3. Although increasing either coverage or heritability results in smaller set sizes, increasing coverage beyond 100 gives diminishing returns as the observed expression levels approach the true expression levels. As expected, AS-Meta tends to struggle at low AS variance, especially apparent at a standard allelic deviation of 0.55, with a mean set size of 78 at best. This may be due to the large majority of samples falling under the threshold for allelic imbalance at 0.65. To verify that PLASMA is calibrated across the full range of conditions, Figure S4 shows that the 95% credible set sizes have at least a 95% chance of including the causal variant.

**Figure 3:**
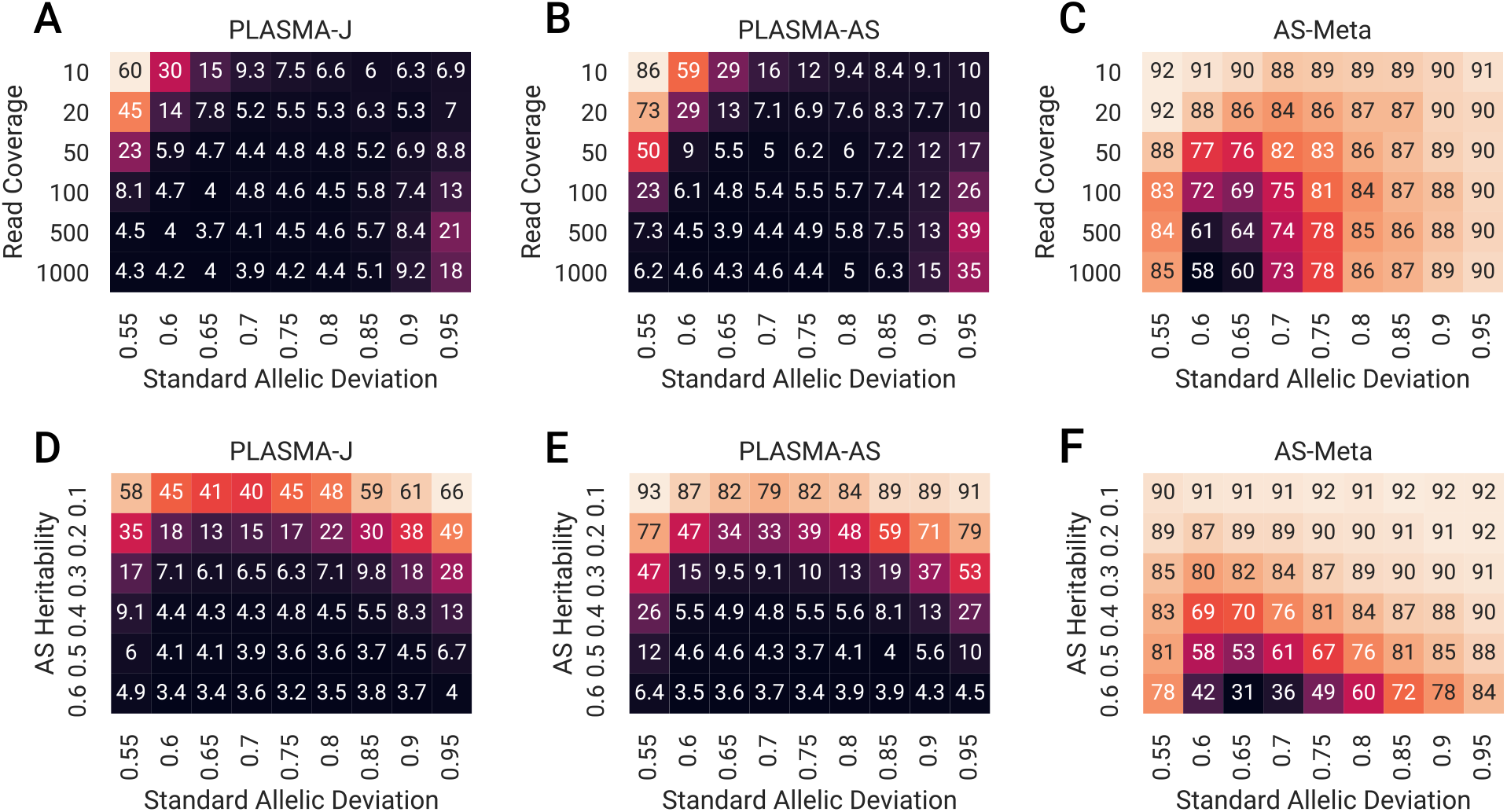
Comparison of the mean 95% confidence credible set sizes across different AS fine-mapping methods. Each square is the mean set size calculated over 500 simulated loci of 100 markers and 100 samples each, with one causal variant. (a, b, c) Mean credible set sizes as a function of standard allelic deviation and mean read coverage, with AS heritability set to 0.4. (d, e, f) Mean credible set sizes as a function of standard allelic deviation and AS heritability, with mean read coverage set to 100 reads.

### 3.3 Inference of multiple causal variants

To demonstrate PLASMA beyond a one-causal-variant assumption, we fine-mapped sets of simulated loci with 2 causal variants with each version of PLASMA. Figure 4a shows the inclusion curves for each version of PLASMA along with FINEMAP. For these curves, inclusion is defined as the expected proportion of causal variants selected, where an inclusion of 1.0 indicates the identification of both causal variants. Here, PLASMA-J and PLASMA-AS deliver an improvement over conventional QTL fine-mapping. Compared to single causal variant fine-mapping, all methods display a decrease in power, which is consistent with results in earlier QTL fine-mapping analysis [13, 14], where capturing all causal variants becomes increasingly difficult as the number of causal variants increase. The lower power for fine-mapping multiple causal variants may be due to the stringent requirement that a model must identify all causal variants in a locus for an inclusion of 1.0. To evaluate the ability of the models to detect the top causal variant, we relax this requirement from identifying all causal variants per locus to at least one causal variant per locus. Inclusion plots for this scenario are shown in Figure 4b, with PLASMA greatly improving the prioritization of the lead causal variant over existing methods.

**Figure 4:**
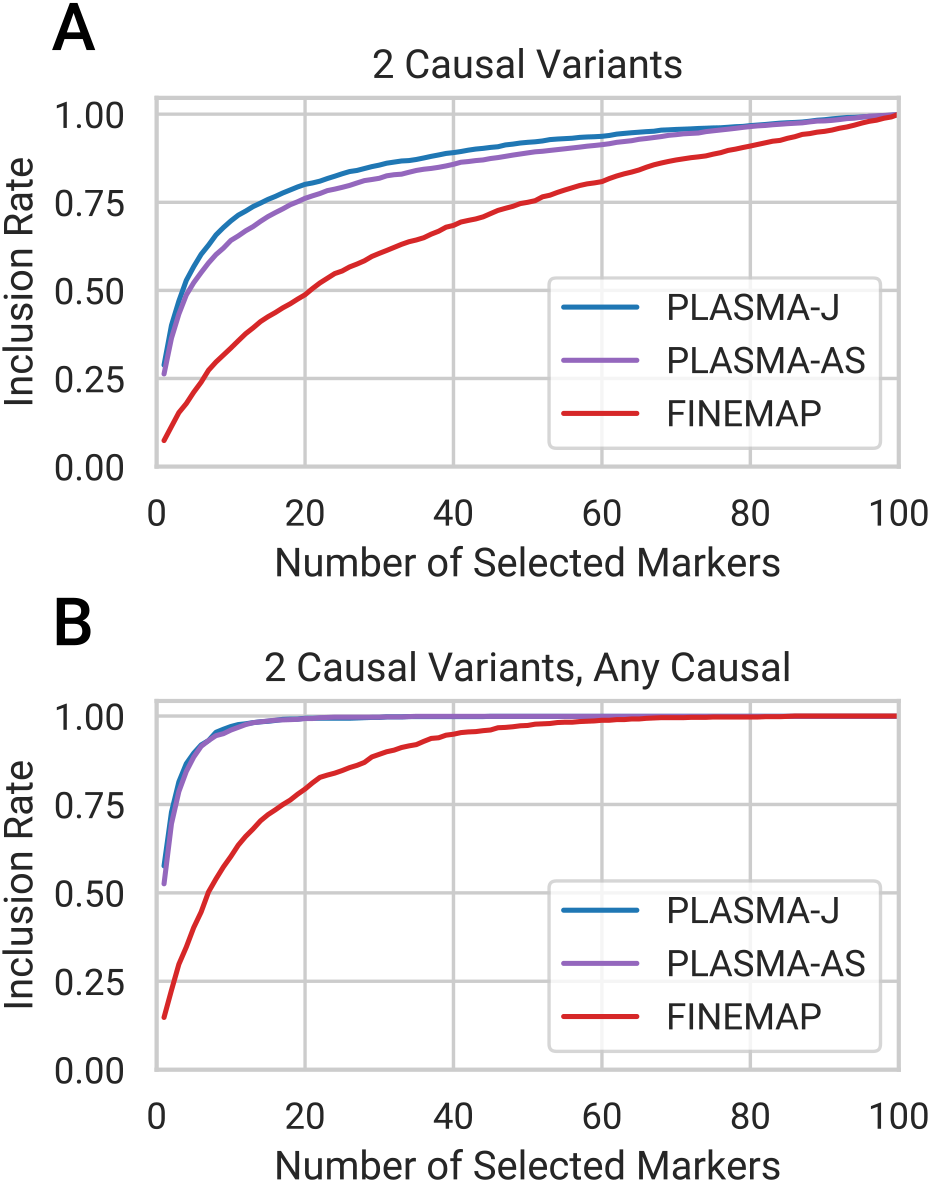
Comparison of fine-mapping methods on sets of simulated loci with two causal variants. Each locus is of 100 markers and 100 samples each, with AS heritability set to 0.4, QTL heritability set to 0.05, and mean read coverage set to 100 reads. (a) Distribution of 95% confidence credible set sizes. (b) Inclusion curve for including both causal variants. (c) Inclusion curve for including one or both causal variants.

We next considered credible set sizes which, unlike the inclusion curves, require accurate calibration to be comparable. Previous analyses have shown that proper calibration of fine-mapping methods is more challenging in the presence of multiple causal variants [30]. Unlike the single causal variant case, where all PLASMA model hyperparameters were inferred from simulation parameters, the causal variance hyperparameters in this case were manually calibrated. This need for calibration may be due to linkage disequilibrium obfuscating the relationship between causal effect sizes and total heritability at a locus, and further complicated by the imperfect estimation of linkage disequilibrium at 100 samples [31]. (See Appendix for information about hyperparameter estimation.) The PLASMA results shown in this section are calibrated such that the recall rates for 95% confidence credible sets are 0.95, 0.96 for PLASMA-J and PLASMA-AS, respectively. This calibration yields median credible set sizes of 86.0 and 90.0 for PLASMA-J and PLASMA-AS, respectively. Like PLASMA, FINEMAP requires user-defined hyperparameters on the prior number of causal variants and on the causal effect sizes. These FINEMAP parameters were set to be equivalent to corresponding calibrated PLASMA parameters. Despite this conservative parameter setting, FINEMAP is overconfident on this dataset with a recall rate of 0.86, so the generated credible sets for FINEMAP are not directly comparable to those of PLASMA.

### 3.4 Fine-mapping of TCGA kidney RNA-Seq data

To evaluate our method on real data, we fine-mapped gene expression data from 524 human kidney tumor samples and 70 matched normal samples collected by TCGA [32]. The data was processed through a rigorous QC pipeline to account for mapping biases based on established best practices [19, 22]. Figures 5a and c show credible set size distribution plots for tumor and normal data, respectively, under a 1 causal variant assumption. We furthermore analyze how well often each method is able to narrow down credible sets under a certain size in simulated loci with one causal variant, shown in Figures 5b and d. Among the tumor samples (N=524, 5652 loci), 27.9% of loci are fine-mapped within 10 variants with PLASMA-J, while 3.4% of loci are fine-mapped within 10 variants with FINEMAP. Furthermore, 263 of these loci can be fine-mapped down to a single causal variant by PLASMA-J. PLASMA-J moreover achieves a median credible set size for 32 variants, whereas FINEMAP achieves a median credible set size of 167 variants. FINEMAP also significantly improves over AS-Meta, which has 6.6% of loci fine-mapped within 10 causal variants, and a median credible set size of 166. Results for normal samples (N=70, 2034 loci) have a similar trend, with 23.2%, 2.5%, and 1.3% of loci fine-mapped within 10 causal variants, for PLASMA-J, AS-Meta, and FINEMAP respectively. Corresponding median credible set sizes are 32, 248, and 252 variants, for PLASMA-J, AS-Meta, and FINEMAP respectively. The somewhat lower power for all models is due to having fewer normal samples than tumor samples. To show that these credible set sizes are robust our choice of heritability hyperparameters, fine-mapping analyses were repeated on the full set of tumor genes with the AS heritability hyperparameter set to 0.05 instead of 0.5. A comparison of the credible set sizes with those from the original parameters are shown in Figure S6.

**Figure 5:**
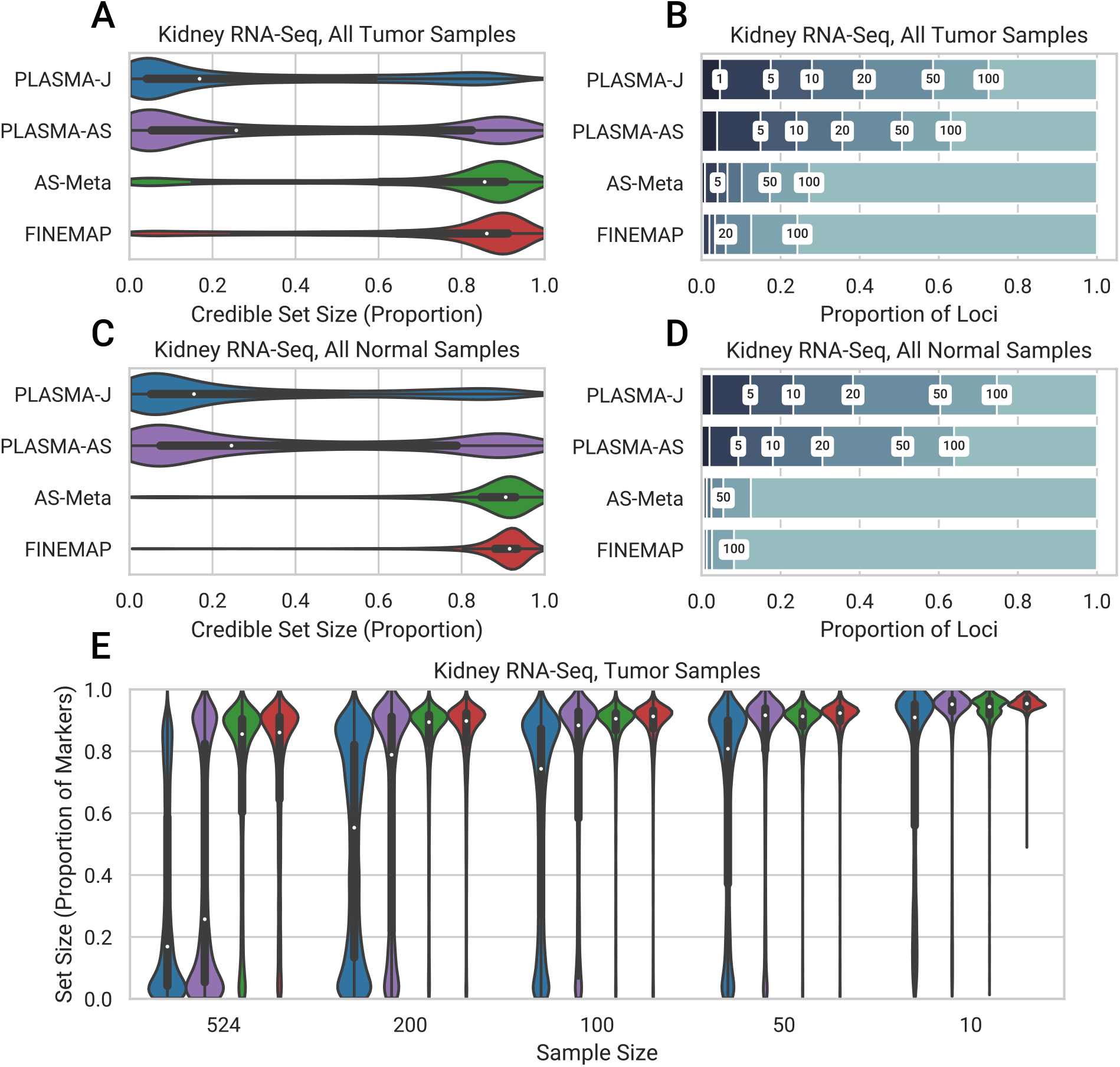
Comparison of the distribution of 95% confidence credible set sizes across loci in kidney tumor and normal samples, with an allelic imbalance false discovery rate of 0.05. Expression was measured as RNA-Seq read counts mapped to phased genotypes. Fine-mapping was conducted with 500 tumor samples and 70 normal samples. Every SNP within 100 kb of each locus was included in the fine-mapping input. (a, c) Distribution of credible set sizes for tumor and normal samples, respectively, under a 1-causal-variant assumption. (b, d) Proportion of loci fine-mapped to 95% credible sets within given thresholds. Thresholds used are 1, 5, 10, 20, 50, and 100. (e) 95% credible set size distributions for randomly down-sampled kidney tumor datasets with decreasing sample sizes.

To investigate how the methods perform at lower sample sizes, we randomly subsample individuals prior to fine-mapping. Figure 5e plots the credible set size distributions for PLASMA-J, AS-Meta, and FINEMAP at various sample sizes of kidney tumor data. In terms of loci fine-mapped to credible set sizes within 10 variants, PLASMA with 50 samples (484 loci within 10 causal variants) has significantly greater power than FINEMAP with 500 samples (193 loci within 10 causal variants) or AS-Meta with 500 samples (371 loci within 10 causal variants). Additionally, in terms of median credible set size, PLASMA with 10 samples (170 median) has about the same power as FINEMAP with 500 samples (167 median). At a given sample size, PLASMA is thus better able to prioritize variants that will be ranked highly in larger studies. Furthermore, as sample size increases, PLASMA increases in power relative to other methods. In tumor samples PLASMA yields a 1.3-fold decrease in median credible set size over FINEMAP at 10 samples, but a 6.9-fold decrease at 500 samples, indicating that PLASMA scales more effectively with sample size than conventional QTL fine-mapping. Nevertheless, PLASMA yields a substantial reduction of credible set sizes even with sample sizes as low as 10, with a median credible set size of 170, compared to a median set size of 219 with FINEMAP. An analogous down-sampling analysis on the normal data is shown in Figure S7. There, PLASMA has higher power for normal samples than for tumor samples, which may be due to the lower variance in the normal data.

Next, we look at how causal variant prioritization is impacted by sample size in the down-sampled analysis. Because the true causal variants in each locus is not known, we use a proxy of markers with a posterior probability of at least 0.1 when fine-mapped with FINEMAP on all samples. Note that this will strongly bias the credible set in favor of FINEMAP and thus do not compare this proxy to FINEMAP credible sets. In Figure S8, PLASMA is again more effective than AS-Meta at each sample size at prioritizing causal variants.

Lastly, to explore multiple causal variant fine-mapping on real data, we run PLASMA and FINEMAP assuming up to three causal variants on the full tumor and normal kidney RNA-Seq dataset. Figure S9 shows multiple causal variant fine-mapping results for kidney tumor and normal RNA-Seq data. As with the simulations, all methods increase in credible set sizes relative to single-causal-variant fine-mapping. On tumor data, PLASMA-J, PLASMA-AS, and FINEMAP report a median credible set size of 93, 172, and 150 respectively, with the caveat of possibly unstable calibration for multiple causal variants (as seen in simulations). Interestingly, PLASMA-AS displays a larger power drop than FINEMAP does. This difference suggest that allelic imbalance may be less informative when fine-mapping with multiple causal variants. Nevertheless, PLASMA-J performs substantially better than either, suggesting that the joint model is able to combine power from both QTL and AS signals. Regardless, it appears that the majority of loci contain a single causal variant, with FINEMAP estimating this fraction at 68.8%.

Lastly, we look at how PLASMA prioritizes experimentally-verified causal variants at GWAS risk loci. Figure 6 shows the strength AS and QTL associations for DPF3 and SCARB1, genes in two kidney GWAS loci that have verified causal variants [23, 33]. At each sample size threshold, the AS statistic more confidently identifies the true causal variant than the QTL statistic. In the case of DPF3, the AS statistic is able to prioritize the true causal variant at a substantially lower sample size than the QTL statistic. Moreover, the 95% credible sets from the PLASMA-AS model are smaller than those from the QTL-Only model at a given sample size. By producing a more accurate and confident prioritization of causal variants, PLASMA can substantially reduce the difficulty of experimentally validating causal variants.

**Figure 6:**
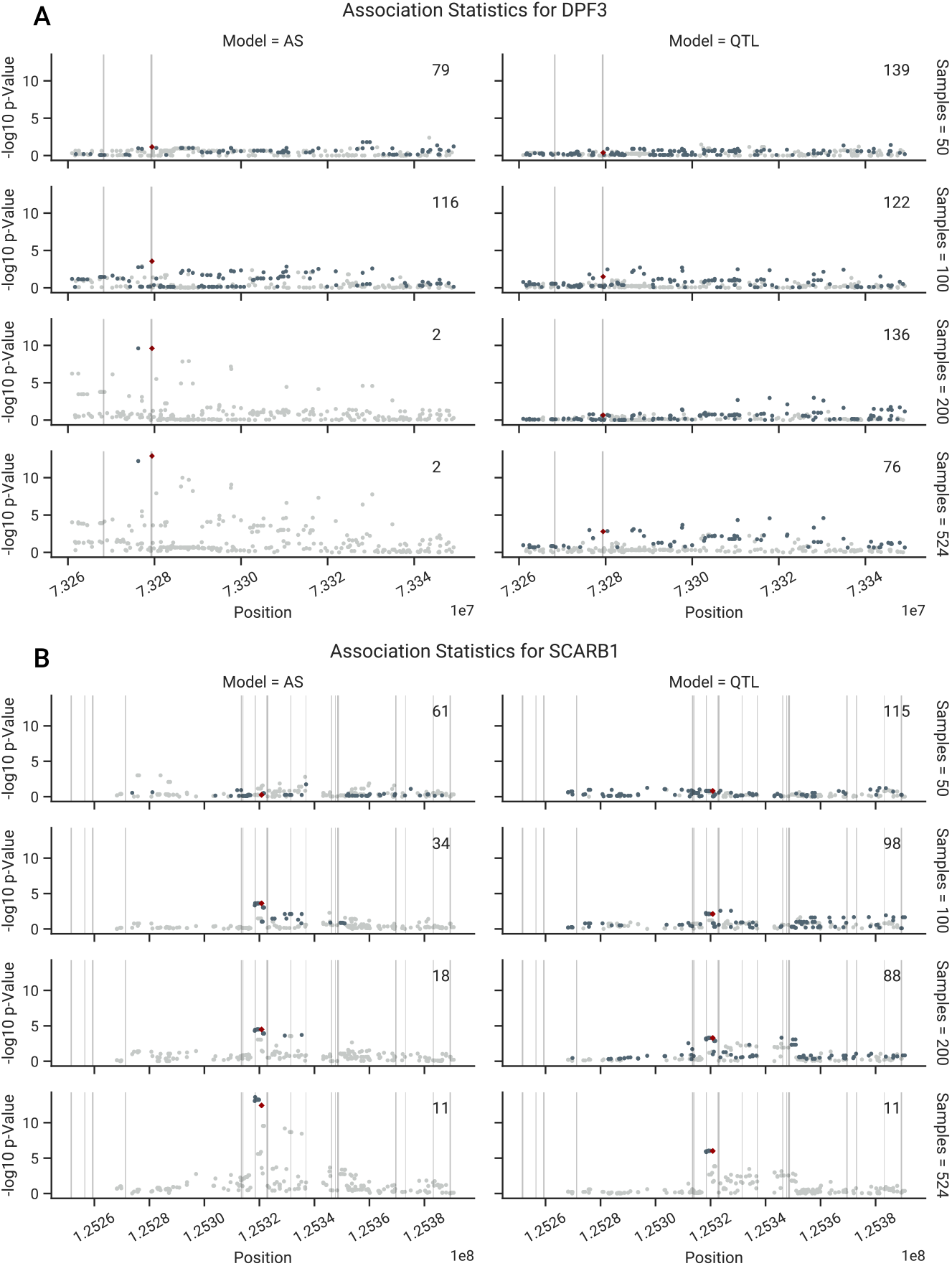
Comparison of AS and QTL associations in the experimentally-verified loci, as a function of sample size. Regions of 70 kb are shown around the causal marker. AS and QTL p-Values were calculated using from *z_ϕ_* and *z_β_*, respectively. Fine-mapping was conducted with the PLASMA-AS and QTL-Only models, respectively. 95% credible set sizes for the whole locus are displayed in the top right of each subplot. Markers in the 95% credible set are shown in dark gray, while markers not in the sets are shown in light gray. The experimentally verified causal marker is shown in red. Regions of open chromatin (DNAse-Seq peaks) are shaded in gray. (a) Associations in the DPF3 locus (b) Associations in the SCARB1 locus.

### 3.5 Fine-mapping of prostate H3k27ac ChIP-Seq data

To evaluate PLASMA with a different molecular phenotype, we fine-mapped H3k27ac activity measured by ChIP-seq from 24 human prostate tumor samples and 24 matched normals. Although this study measures chromatin activity rather than expression, the nature of the data is nearly identical to that of RNA-Seq and is processed analogously by our QC pipeline and by PLASMA. Instead of fine-mapping eQTLs around gene loci, we fine-mapped chromatin QTLs (cQTLs) around chromatin peaks. Figures 7 shows distribution plots for tumor data (1375 peaks) and normal data (908 peaks) under a 1 causal variant assumption. Among the tumor data, 14.5% of peaks are fine-mapped within 50 variants with PLASMA-J, while 1.9% of loci are fine-mapped within 50 variants with FINEMAP. Furthermore, PLASMA achieves a median credible set size of 236, compared to QTL-Only fine-mapping achieving a size of 318. PLASMA also outperforms AS-Meta, with 1.9% of loci fine-mapped within 50 causal variants (no gain over FINEMAP), and a median credible set size of 310. Results from normal samples are comparable, with 5.2%, 2.5%, and 2.3% of loci fine-mapped within 50 causal variants, for PLASMA-J, AS-Meta, and FINEMAP respectively. These methods achieve a median credible size of 296, 313, and 319 variants, respectively. Overall, these ChIP fine-mapping results are roughly in line with those from RNA-Seq fine-mapping.

**Figure 7:**
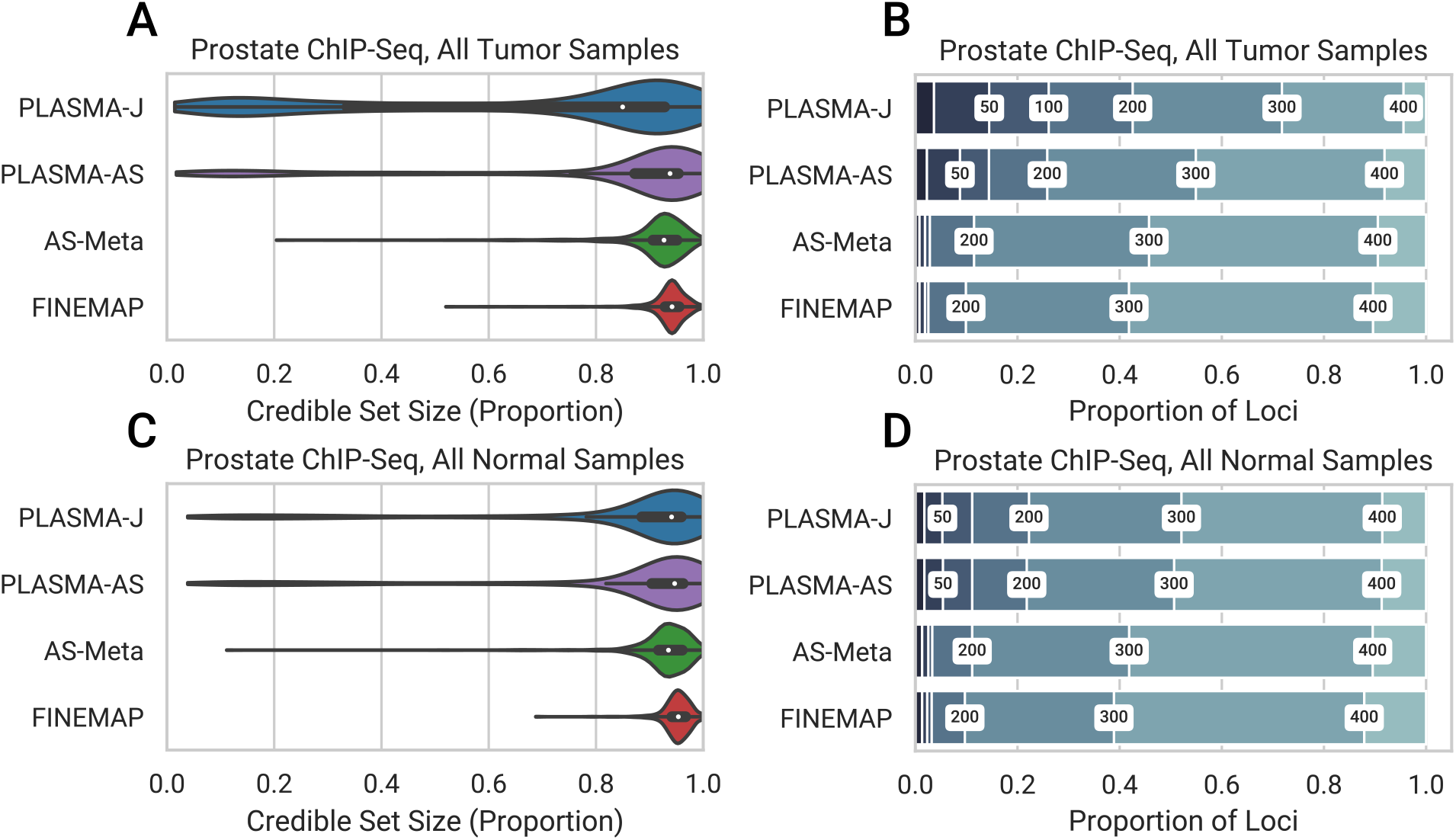
Comparison of 95% confidence credible set sizes for peaks in prostate tumor and normal cells, with an allelic imbalance false discovery rate of 0.05. Presence of H3k27ac histone marks was quantified as ChIP-Seq read counts mapped to phased genotypes. (a, c) Distribution of credible set sizes for tumor and normal samples, respectively, under a 1-causal-variant assumption. (b, d) Proportion of peaks fine-mapped to 95% credible sets within given thresholds. Thresholds used were 20, 50, 100, 200, 300, and 400.

### 3.6 PLASMA increases functional enrichment of credible set markers

To evaluate PLASMA’s ability to select markers in functional regions using kidney RNA-Seq data, we look for enrichment of prioritized variants at open chromatin regions measured with DNAse-Seq in a kidney cell-line [34]. Since chromatin accessibility is an indicator of transcription factor binding and regulation [35], an enrichment of credible set markers for open chromatin would indicate that the fine-mapping procedure is prioritizing markers in functionally relevant regions. For instance, the causal variant in the DPF3 locus lies within a DNAse-Seq peak (Figure 6a). Note that quantifying overlapping with an independent functional feature such as open chromatin imposes no assumptions on the ground truth, in contrast to comparing to external QTL/GWAS data which may be biased towards conventional QTL analysis. The null distribution is defined as the credible set markers being located independently of open chromatin and use Fisher’s exact test to calculate enrichment as a function of minimum causal variant probability. Figures 8, S10a and S10c,d show the odds ratios and p-values (computed by Fisher’s exact test), respectively, as a function of posterior probability threshold from each fine-mapping method. The credible set markers produced by PLASMA, for the most part, display a significantly stronger enrichment with open chromatin compared to existing methods. For instance, at the *p* = 0.1 threshold for tumor samples, PLASMA’s credible set markers achieve a p-value of 9.26 ×10^−52^ and an odds ratio of 2.16. In comparison, credible sets from QTL-Only fine-mapping at that threshold achieves a p-value of 2.02 ×10^−7^ and an odds of 1.62. This enrichment shows that even with far smaller credible sets, PLASMA is able prioritize markers that fall in regions of likely functional significance. The difference between PLASMA and existing methods is greatest at higher posterior probability thresholds. PLASMA may be assigning a more meaningful number of markers with such high posterior probabilities, compared to existing methods that are rarely so confident about a marker’s causal status.

**Figure 8:**
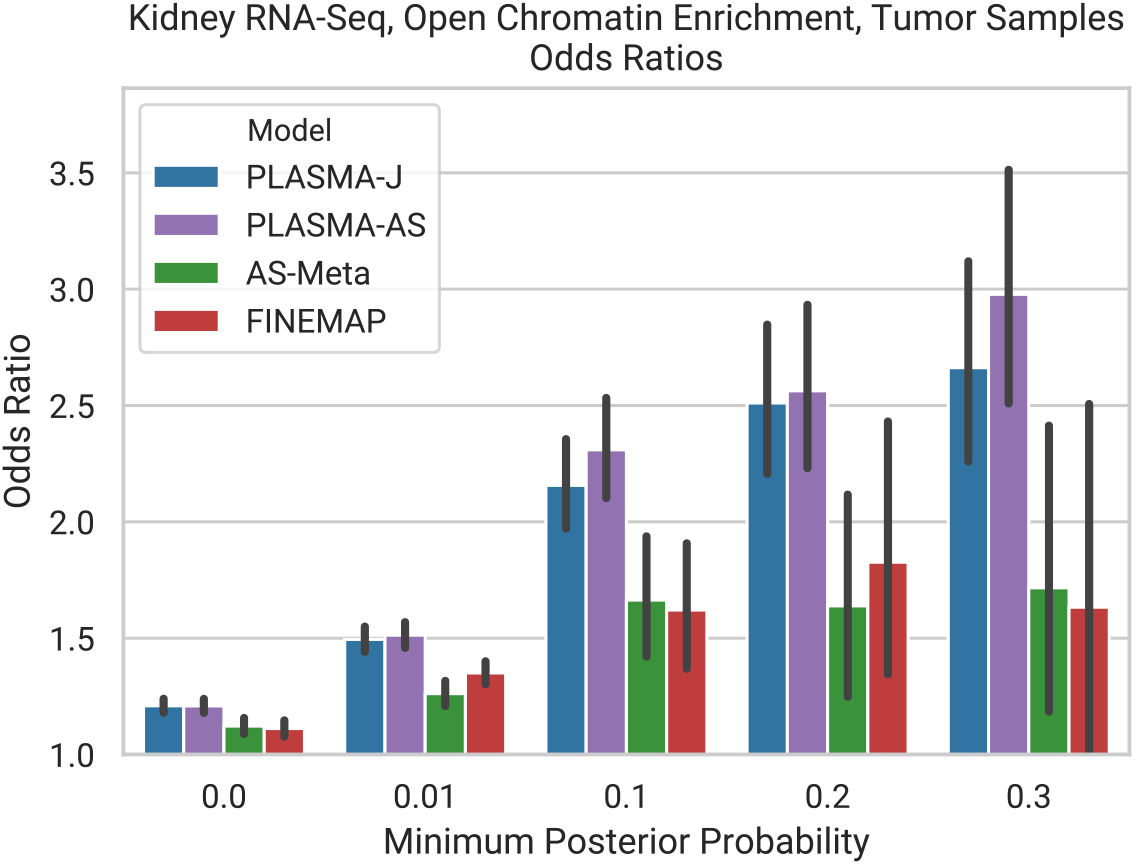
Enrichment of PLASMA kidney RNA-Seq credible sets on open chromatin. The x-axis represents a minimum posterior probability threshold on the markers within credible sets, with a higher threshold indicating more confident but fewer markers. The markers in these thresholded credible sets were intersected with the functional feature in question. 95% confidence intervals were calculated with Fisher’s exact test, with a control of all markers in a locus.

Similarly, to validate the credible sets computed from prostate ChIP-Seq data, we look for enrichment of credible set markers at chromatin looping anchors measured by Hi-ChIP in a prostate cell-line. Regulatory elements overlapping loops are more likely to be involved in cis-regulation and we reasoned that they should therefore be enriched for true causal cQTLs [36, 37]. Again, we note that this functional feature is independent of the QTL signal or locus LD and is not biased towards a QTL or AS model. Figure S10b and S10e show the odds ratios and p-values, respectively, across models as a function of posterior probability threshold (computed by Fisher’s exact test). The credible set markers produced by PLASMA display a significantly stronger enrichment with looping anchors compared to the other methods. For instance, at the *p* = 0.1 threshold, PLASMA’s credible sets achieve a p-value of 1.05 × 10^−6^ and an odds of 1.77. In contrast, credible set markers from FINEMAP at that threshold achieves a non-significant p-value of 0.80 and an odds of 0.72.

## 4 Discussion

We present PLASMA, a statistical fine-mapping method that utilizes allele-specific expression and phased genotypes to select candidate causal variants. By modeling gene expression at a locus in an allele-specific manner, PLASMA scales in power both across individuals and across read counts. Through read-count-level simulations of loci, we show that PLASMA performs robustly across a wide range of realistic conditions and consistently outperforms existing statistical fine-mapping methods, including cases where a significant amount of observed imbalance is caused by non-genetic factors. We further demonstrate this increased power on experimental data by applying PLASMA to a large RNA-Seq study, as well as a smaller ChIP-Seq study. In both cases, PLASMA achieves substantially smaller credible set sizes compared to existing fine-mapping methods, greatly increasing the number of loci amenable to experimental causal variant validation. Lastly, we show that even with these greatly reduced (more specific) credible set sizes, PLASMA achieves an equivalent or superior degree functional enrichment as existing methods. These results not only present PLASMA as a powerful tool for prioritizing causal variants, but also demonstrate how AS analysis can be directly integrated into statistical fine-mapping. A key benefit of PLASMA is its ability to utilize existing, conventional sequencing-based QTL data, such as RNA-Seq, CHiP-Seq, and ATAC-Seq at low sample size. This allows researchers to gain significant insight simply by revisiting past QTL studies, especially those with sample sizes too low for conventional QTL fine-mapping.

Although it is evident that an AS analysis with PLASMA confers more signal than an equivalently-sized QTL analysis, AS analysis presents additional obstacles and potential confounders. First, unlike conventional QTL fine-mapping methods that rely only on allelic dosage, PLASMA additionally utilizes genotype phasing, making phasing accuracy a potential concern. However, since PLASMA focuses on cis-regulation, the genotypes observed span no more than several hundred kilobases per locus, well within the high accuracy range of modern phasing algorithms [38]. Second, PLASMA depends on having heterozygous individuals in the tested feature and SNP in order to leverage AS signal. In our analyses we focused on features that were testable by AS (10946 of 19645 total genes, 113459 of 525629 total peaks). However, even in the complete absence of heterozygotes, PLASMA can still conduct conventional fine-mapping based on dosage and total expression. Recent technologies that could potentially offer greater signal include RNA-seq with unspliced transcripts [39], and direct allele-specific measurement of expression using single-cell RNA-Seq [40]. Third, PLASMA assumes the same causal configuration underlying both the AS and QTL effects (and is thus able to combine the signals) but the causal effects may differ due to real biological confounding. For example, cis effects on gene A followed by (local) trans effects of gene A on gene B would be identified as a QTL association, but would not exhibit AS association. This would be a model violation for PLASMA and produce larger credible set sizes. Although PLASMA can consider correlations between causal AS and QTL affect sizes, this parameter is hard to estimate, and we find in real data that the model with correlation set to zero (PLASMA-J) exhibited greater power than a nonzero constant. Future work is required to fully elucidate the relationship between allele-specific and total effects, which likely differs across genes. Fourth, genomic imprinting (where either the maternal or paternal copy of the gene is silenced) or random monoallelic expression would produce the appearance of allelic imbalance within affected individuals in the absence of true cis-regulatory signal [20]. Although PLASMA does not explicitly model such biases, a bias that is independent of genotype will only cause a reduction in power and not produce false-positives. A potential extension would be to model such violations or discrepancies between the QTL and AS models directly, following the lines of methods such as RASQUAL [20]. Fifth, PLASMA currently does not incorporate covariate analysis in the allele-specific model (though the intra-individual nature of the test controls for false positives), which could additionally be used to model environmental confounders and increase power [41]. AS covariate analysis could potentially be achieved through a multivariate likelihood ratio test as in WASP [19].

PLASMA’s approach in combining QTL and AS signals opens up possible future work in two distinct directions. The first direction would be to build upon the generative fine-mapping model to incorporate additional sources of signal. For example, one can incorporate epigenomic annotation data by setting the priors for causality for each marker. Approaches used in existing QTL-based methods such as PAINTOR and RiVIERA-MT [16, 42] could be transferred to PLASMA with relatively little difficulty. Another possibility would be to conduct *N*-phenotype colocalization by utilizing additional phenotypes in addition to the AS and QTL phenotypes. Generalizing from two to multiple phenotypes would be straightforward, and could utilize the colocalization algorithm first introduced in eCAVIAR [2]. A second, more general direction would be to adapt QTL-based population genetics methods to utilize AS summary statistics. Since both QTL and AS statistics can be characterized as linear combinations of haplotype-level genotypes, they share many distributional properties, including LD, allowing them to be easily interchangeable in many circumstances. One such application would be gene expression prediction for transcriptome-wide association studies (TWAS) [43], where the increased signal of AS statistics could increase power to identify gene-phenotype relationships. Broadly speaking, the allele-specific model and association statistics that PLASMA introduces will be relevant to any analysis of small sample size or limited tissue.

## Acknowledgments

We thank F. Hormozdiari for guidance on statistical fine-mapping and C. Kalita for guidance on model validation. We also thank B. Pasanuic and C. Giambartolomei for helpful feedback.

## 5 Appendix

### 5.1 Modeling total expression at a locus (QTL)

#### 5.1.1 Modeling genetic effects on total expression

We calculate marginal effect sizes for a given locus under the conventional linear model of total gene expression. Let us consider a QTL study of a given locus with *n* individuals and *m* markers. Let **y** be an (*n* ×1) vector of total expression across the individuals, recentered at zero. Given a marker *i*, let **x***_i_* be an (*n* × 1) zero-recentered vector of genotypes. We define *β_i_*, the genetic effect of marker *i* on total gene expression as follows:

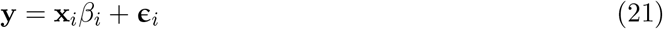

We model the residuals **ϵ***_i_* as normally distributed with variance 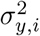.

#### 5.1.2 Calculation of QTL summary statistics

We use the maximum likelihood estimator of *β_i_*, equivalent to the ordinary-least-squares linear regression estimator:

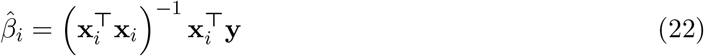

Under the null model where *i* is not causal, *i* does not explain any amount of variation of the phenotype, and the variance of **y** is simply 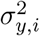. Thus, under the null:

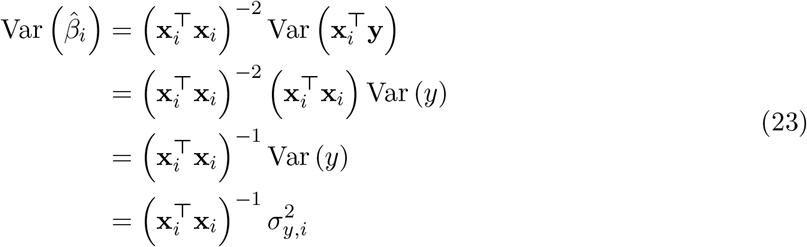

We estimate 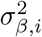 from the residuals:

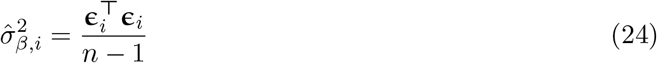

We thus define our QTL summary statistic (Wald statistic) for marker *i* as:

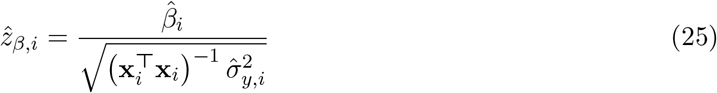

We assume that the number of individuals is enough such that the observed statistic is normally distributed with unit variance:

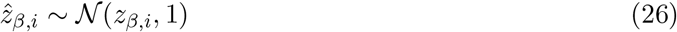

In the case where **x**_*i*_ is of unit variance, the statistic simplifies to:

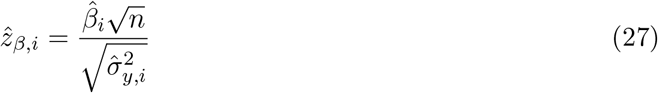

### 5.2 Modeling allele-specific expression at a locus (AS)

#### 5.2.1 Modeling haplotype-specific effects on expression

We model allele-specific expression under the observation that a cis-regulatory variant often has a greater influence on the gene allele of the same haplotype. Under this model, an individual who is heterozygous for one or more cis-regulatory markers will show an imbalance in expression between the alleles.

From a quantitative perspective, let us consider a single locus in a single individual who is heterozygous for marker *i*. Let 0 and 1 represent the wild-type and alternative marker alleles, respectively. We define *e*_0_ as the expression of the gene allele on the same phase as marker allele 0, and *e*_1_ as the expression of the gene allele on the same phase as marker allele 1. Let 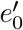 and 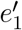 be baseline expressions without the effect of marker *i*. We define *δ_i_* as the cis-regulatory strength of marker allele 1 over marker allele 0 such that:

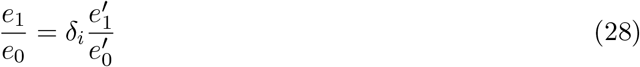

If we define *i*’s phase, *v_i_*, we can arbitrarily assign haplotypes *A* and *B*. The above equation then becomes:

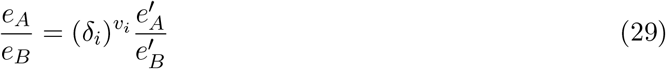

The marker’s phase is 1 if haplotype *A* contains the alternative marker allele, −1 if haplotype *B* contains the alternative marker allele, and 0 if the individual is homozygous for the marker.

We now re-write Equation 29 as a linear model. Let *w* be the log expression ratio between haplotypes A and B:

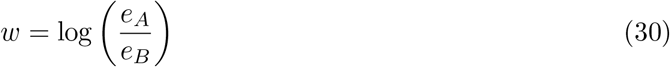

Let *ϕ_i_* be the log allelic fold change (logAFC) caused by variant *i*:

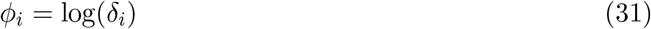

Let *ζ_i_* be the log baseline expression ratio between haplotypes A and B:

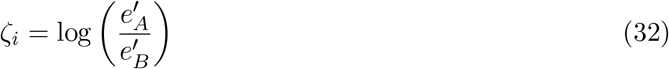

With these parameters we rewrite Equation 29 as:

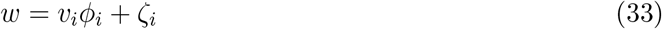

Given *n* individuals, this expression becomes:

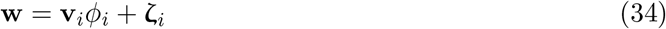

We assume that ζ*_i_* is drawn from a normal distribution with variance 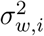. Note that under this model, *ϕ_i_* can be interpreted as the effect size of marker *i* on allelic imbalance, with ζ*_i_* as the residuals. Furthermore, assuming no haplotype bias, both **w** and **v**_*i*_ are zero-centered in expectation.

Experimentally-derived AS data, such as RNA-Seq data, yield reads that are mapped to a particular haplotype. Given *c_A_* and *c_B_*, the read counts mapped to haplotypes *A* and *B* respectively, we define our estimator of *w* as:

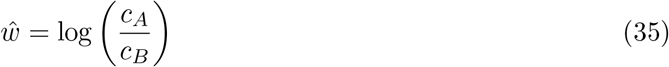

For a given individual *j*, we define *c_A,j_* as the allele-specific read count from haplotype *A*. We model the allele-specific read count as drawn a beta-binomial distribution, given the total mapped read count *c_j_*:

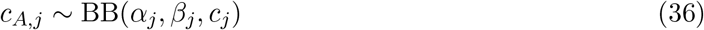

We define *π_j_* as the expected proportion of read counts (allelic fraction) from haplotype *A*:

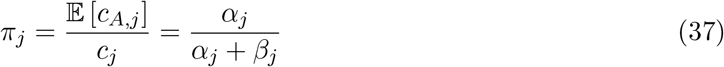

*α_j_* and *β_j_* can be re-parameterized in terms of *π_j_* and the sampling overdispersion *ρ_e_*.

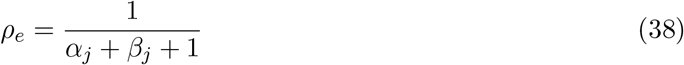

With this re-paramaterization, the mean and variance of *c_A,i_* is given as follow:

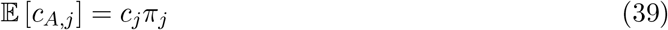

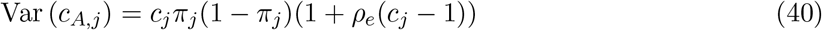

We use this beta binomial model to estimate the variance of *ŵ_i_*. We scale the distribution by 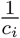 to get the mean and variance for the read count proportion:

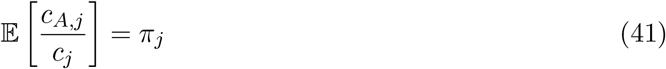

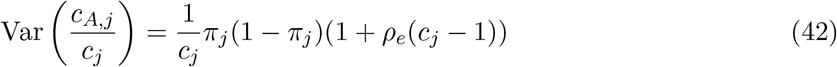

We define *w*\* as the logit-transformed allelic fraction:

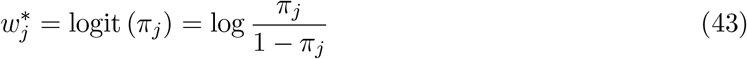

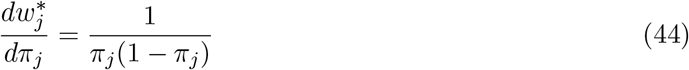

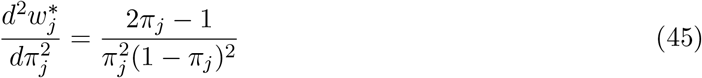

We can thus find the approximate mean and variance of *ŵ_j_* given 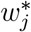 using Taylor expansions:

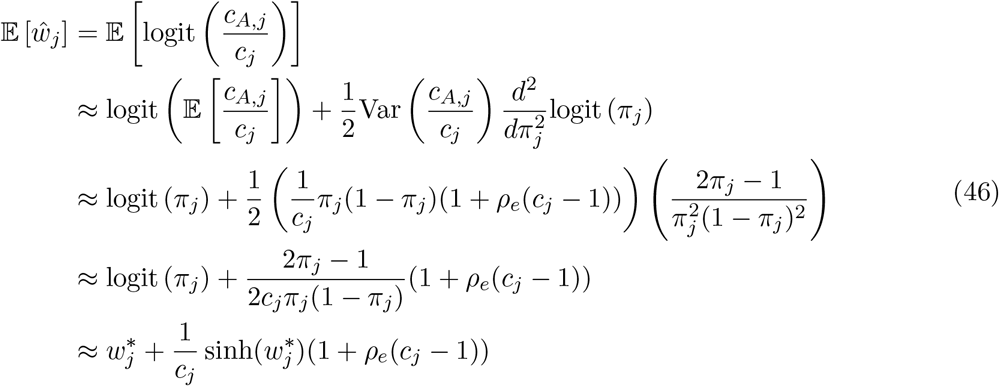

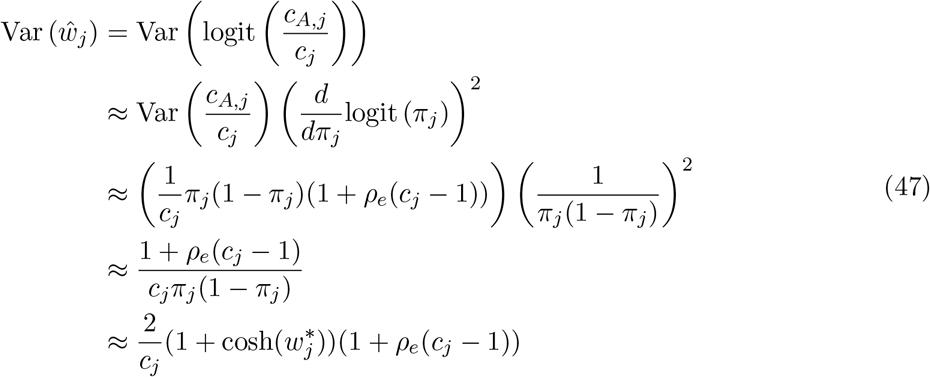

Note that *w* and *w*\* are not equivalent because 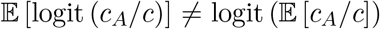. Equation 46 implies that *ŵ* is a biased estimator of *w*\*, especially at low read counts and/or high overdispersion. To get an estimator of *w*\* with reduced bias, we take the approximation that sinh(*w*\*) ≈ *w*\* around zero:

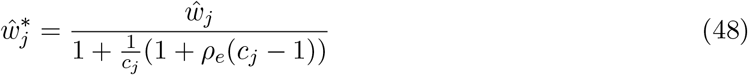

We use *ŵ*\* to find an estimator of 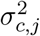, the variance of *ŵ*:

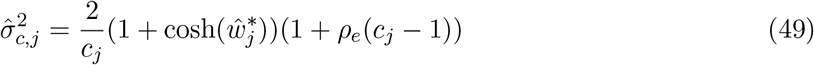

Given our estimator *ŵ_j_*, we quantify the sampling error *τ_j_* = *ŵ_j_* − *w_j_*, with 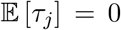 and Var 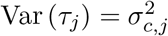. Thus, across individuals:

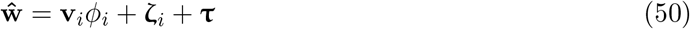

#### 5.2.2 Calculation of AS summary statistics

Due to heteroscedasticity among individuals, we estimate the AS effect size *ϕ_i_* in a weighted manner, giving larger weights to individuals with lower expected sampling error. Given individual *j*, we define the weight for *j* as the inverse of the estimated read count variance:

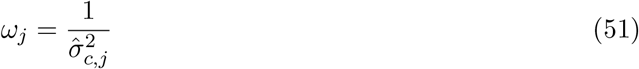

We define our weight matrix **Ω** as a diagonal matrix with **Ω**_*j,j*_ = *ω_j_*.

We use the weighted-least-squares estimator for *ϕ_i_*:

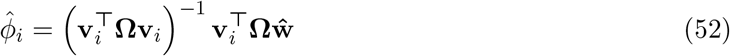

Under the null model where *i* is not causal, the variance of *w_j_* is 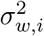, and the variance of *ŵ_j_* is 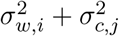. We assume that the experimental errors *τ* and biological residuals *ζ_i_* are uncorrelated. Thus, under the null:

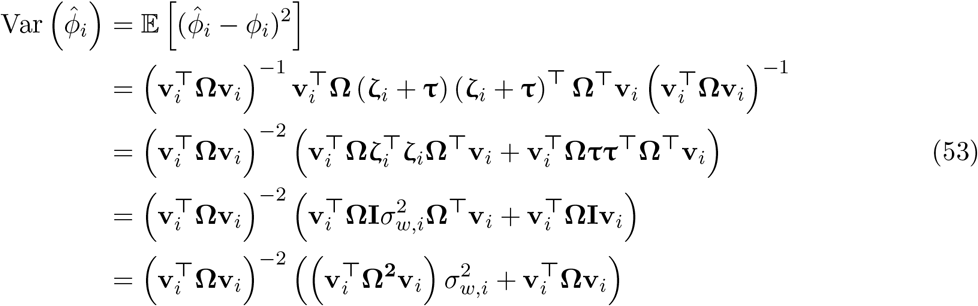

We now estimate 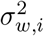 from the residuals. Note that we are estimating the variance of the biological residuals Var (*ζ_i_*), which is distinct from the total residuals are **ζ**_*i*_ + **τ**, so we cannot directly use the variance of the total residuals. We instead use the following estimator for 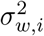:

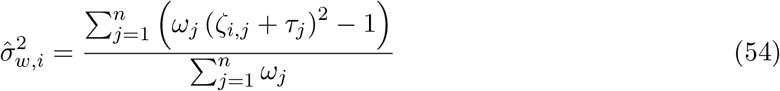

We show that this estimator is equal to 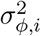 in expectation:

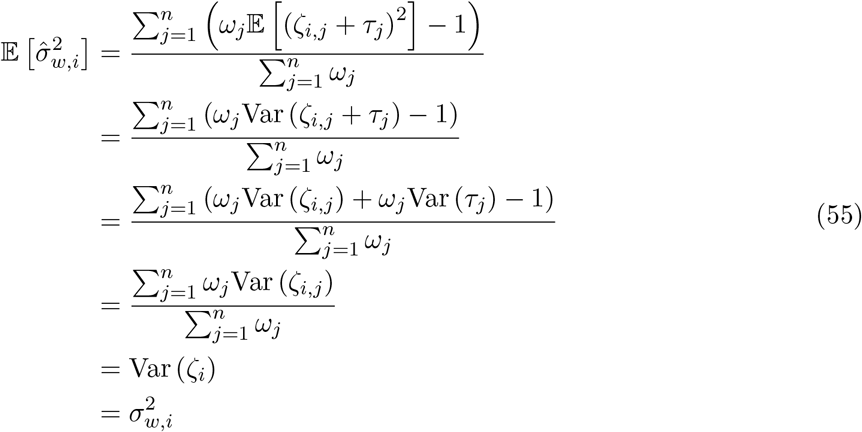

With this estimator, we define the AS association statistic for marker *i* as follows:

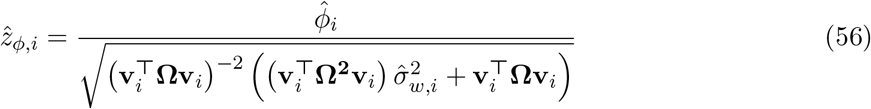

We assume that the observed statistic is normally distributed with unit variance:

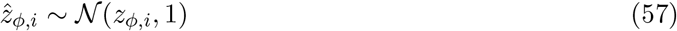

To gain an intuitive understanding of the association statistic, let us examine it under simplifying conditions. We assume that **v**_*i*_ is of unit variance, that read count overdispersion is negligible, and that allelic imbalance and read coverage are fixed across individuals. Under these conditions, let 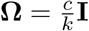 for coverage c and some constant *k*. Equation 56 simplifies to:

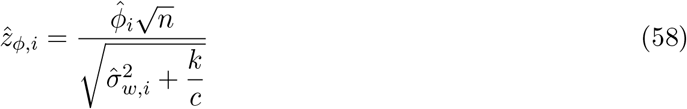

We can see that under high experimental noise (*k/c*), the denominator is dominated by the quality of data (read coverage). In contrast, when experimental noise is low, the denominator is dominated by 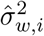, determined by the inherent heritability of the locus’s AS phenotype.

In the case where phasing error is significant, we would expect the estimated AS effects 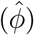 to have more deviation from the true effects. We derive a correction for the AS z-score, given a per-marker probability of mis-phasing *ψ_i_*. We define 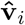 as the imperfect observed phasing for marker *i*, and we define the phasing error vector **δ***_i_* such that 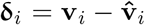. Note that each *δ* is a ternary −2/0/2 indicator, with each *δ*^2^ being a binary 0/4 indicator of a phasing error. We assume that the occurence of a phasing error is independent of which haplotype the alternative allele is one, so that 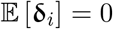. We now derive the variance of 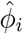 under imperfect phasing:

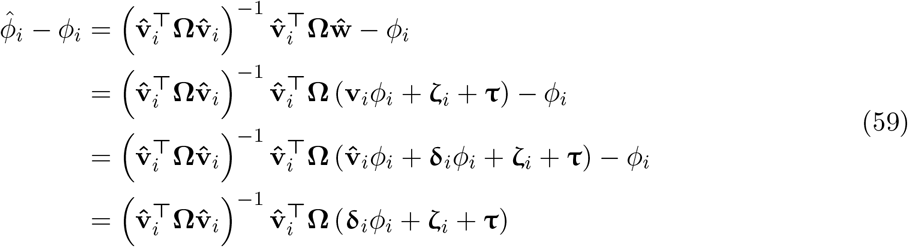

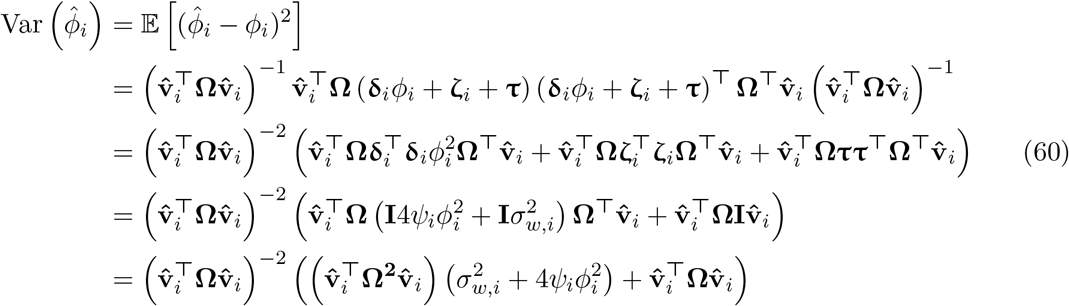

When calculating Var 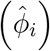, we approximate the 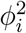 term with the observed 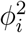. We thus define the corrected z-score:

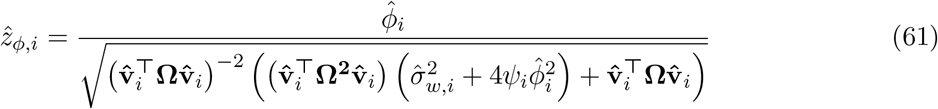

#### 5.2.3 Emprical Analysis of Imperfect Phasing

We evaluate the AS-based methods, PLASMA and RASQUAL+, on simulated loci with imperfect phasing. (We do not include AS-Meta, since the method does not depend on phased genotypes.)We modelled our imperfect phasing simulation on the EAGLE phasing paper [cite], which quantified two sources of phasing error: switch error and blip error. Switch error is the probability that the phasing swaps at a given marker (affecting all downstream markers) and was set to 0.152%. Blip error is the probability that a given marker is misphased, with downstream markers unaffected, and was set to 0.165%.

Results for these imperfectly-phased loci are shown in Figure S5, with comparisons to perfectly phased loci. Comparing the inclusion curves for each method, we see only a very small difference in power with perfect and imperfect phasing, which is expected given the low error rates reported by EAGLE. Similarly, we see a small increase in credible set sizes for each model (RASQUAL+ not included due to non-meaningful sets). However, we do see a modest decrease in recall (0.98 to 0.89 for PLASMA-J, and 0.98 to 0.92 for PLASMA-AS), indicating that imperfect phasing does have some impact on accuracy. Nevertheless, it appears that the overall effect of imperfect phasing is small at real-world error rates.

### 5.3 Inference of causal variants with QTL and AS statistics

#### 5.3.1 Modeling the effects of linkage disequilibrium

PLASMA assumes that the QTL and AS statistics are drawn from the same marker linkage disequilibrium (LD), a property that we verify under the assumption that the two alleles in an individual are distributed independently (See following section for proof). This equivalence yields three LD estimators: using dosage (0/1/2) genotypes *x*, phasing (−1/0/1 genotpyes *v*), or haplotype-specific (0/1) genotypes. To evaluate each of these three estimators and to emprically demonstrate their equivalence in expectation, we sample a contiguous block of 100 SNPs from phased 1000-Genomes genotype data at sample sizes ranging from 50 to 1000. We then calculate LD matrices using each estimator and compare the estimated correlations for each pair of markers. In Figure S1, we see that correlation among the estimators increases as a function of sample size, and also that a sample size of 50 is more than sufficient for high concordance among the three estimators. Since the haplotype-specific estimator appears to be more correlated to both the dosage estimator and the phasing estimator than the latter two are to each other, we use the haplotype-specific estimator in our analyses. We believe that the haplotype-specific estimator is the most accurate of the three since it as effectively double the sample size as the dosage and phasing estimators.

#### 5.3.2 Equivalence in expectation of QTL and AS estimators of LD

Due to linkage disequilibrium, there exist significant correlations of genotypes among markers. This correlation is reflected in the correlations in the association statistics. PLASMA models the QTL (dosage) and AS (phasing) genotypes having an identical LD structure. Here, we show that the LD calculated from the dosage, phasing, and haplotype-specific genotypes are equivalent in expectation, under the assumption that the two alleles in an individual are independent.

Let *a* be a binary genetic marker, and let *p_a_* be a random variable for the (0/1) genotype of a on haplotype I (arbitrarily chosen), and let *q_a_* be a random variable for the (0/1) genotype of a on haplotype II.

We define the dosage genotype of marker a as the sum of the centered haplotypes:

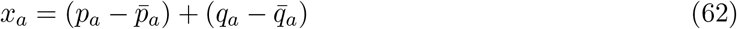

Likewise, we define the phasing genotype of marker *a* as the difference of the centered haplotypes:

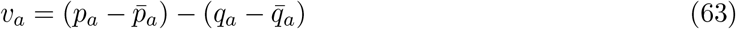

(Note that since 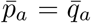, the phasing is also equivalent to the difference of the uncentered haplotypes.)

We now calculate the covariance of *x_a_* and *x_b_* for two markers *a* and *b*. Since these variables are zero-centered, we can express the covariance as 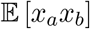:

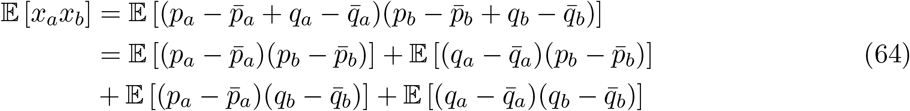

We make the assumption that the two alleles in an individual are independent, so that the covariance of any *p* and *q* is zero. The covariance simplifies to:

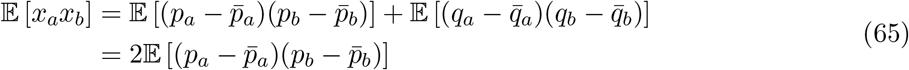

We calculate the correlation of *x_a_* and *x_b_* from this correlation, and we see that it is equivalent to the correlation of *p_a_* and *p_b_*:

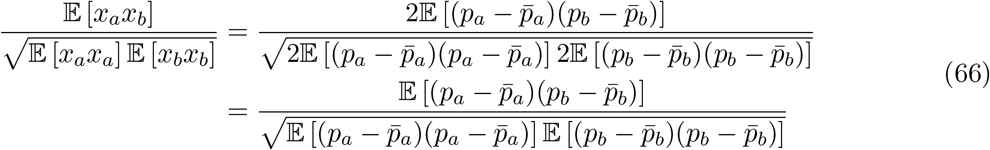

Under the same assumptions, we calculate the covariance of *v_a_* and *v_b_*.

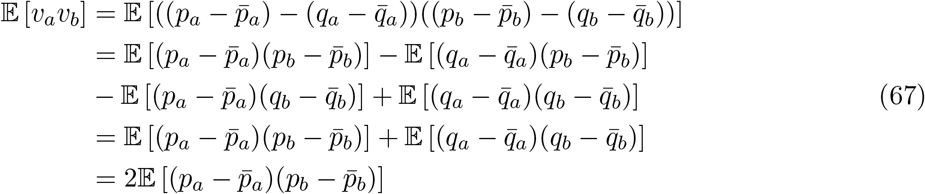

We see that the covariance of *v_a_* and *v_b_* is equal to the covariance of *x_a_* and *x_b_*, implying that the correlation of va and vb is equal to the correlatin of *x_a_* and *x_b_*, and thus *p_a_* and *p_b_*. Therefore, under the assumption that the two alleles in an individual are independent, the LD calculated from dosage genotypes *x*, phasing genotypes *v*, and haplotype-specific genotypes *p* are equivalent in expectation. (Note that this assumption can theoretically be violated if a marker is under selection, but we assume that selection is insignificant due to the weak effects of common markers.)

#### 5.3.3 Jointly modeling total and haplotype-specific effects on expression

We define 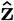 as the combined vector of AS association statistics and QTL association statistics:

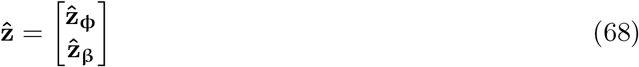

Let *r_βφ_* be the overall correlation between the QTL and AS summary statistics calculated across all loci. We define the combined correlation matrix **R** as:

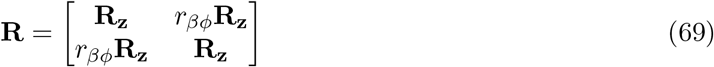

We model the joint distribution as multivariate normal, with covariance **R**:

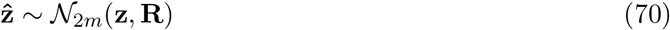

#### 5.3.4 Modeling summary statistics given a causal configuration

The goal of this method is to infer the causal markers, given QTL and AS association statistics. To this end, we introduce a likelihood function that gives the probability of statistics 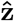, given a causal configuration. We define a causal configuration **c** as a vector of causal statuses corresponding to each marker, with 1 being causal and 0 being non-causal.

Let **z_c,φ_** and **z_c,β_** be the underlying causal AS and QTL effects, respectively, across markers such that:

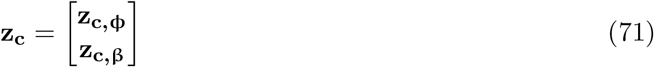

We define hyperparameters 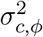 and 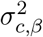 as the variance of AS and QTL causal effect sizes, respectively *r_c,βφ_* as the underlying correlation of the causal QTL and AS effect sizes. (This is not to be confused with *r_βφ_*, which concerns the correlation between the association statistics.) We define **Σ_c_**, the covariance matrix of causal effect sizes, given a causal configuration:

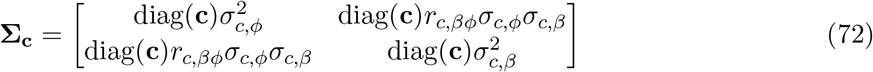

We model the causal effect sizes, given a causal configuration, as drawn from a multivariate normal distribution:

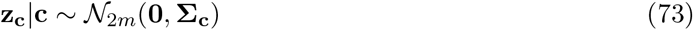

Furthermore, we model the expected association statistic for a given marker as a linear combination of all effects correlated to the marker.

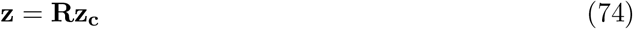

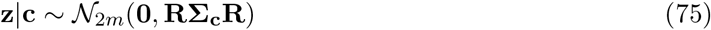

Combining Equations 70 and 75, we get a probability distribution for the observed association statistics given a causal configuration. This is our likelihood for a causal configuration.

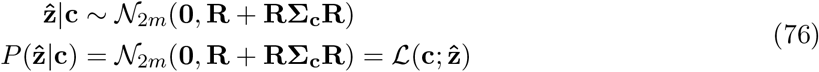

To get a prior distribution for the causal configuration **c**, we define the hyperparameter γ as the prior probability that a single variant is causal and 1 – *γ* as the probability that a variant is not causal. The probability of a configuration consisting of *m* variants thus becomes:

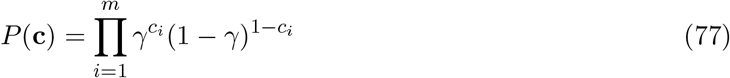

We can view the prior as a regularization term by taking the negative log:

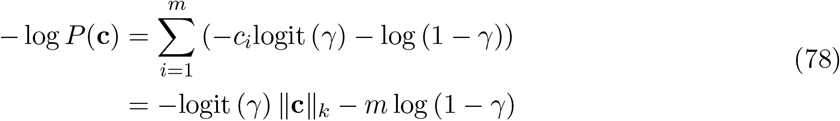

Since **c** is a binary vector, ||**c**||_*k*_ is the same for all positive *k*. Thus, the prior imposes *L_k_* regularization with λ = −logit (*γ*). In practice, this regularization favors causal configurations with fewer causal variants.

With the prior and likelihood, we define the posterior probability of a causal configuration, normalized across the set of all possible configurations 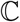:

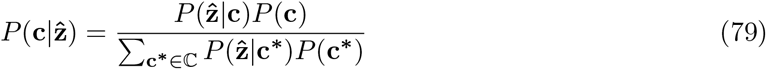

This posterior probability can be alternatively expressed with Bayes Factors. We define the null model as the scenario where all markers are non-causal, so that **c = 0.** The Bayes Factor for a particular **c** would thus be:

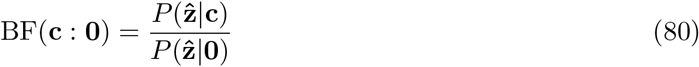

We rewrite Equation 79 with Bayes Factors:

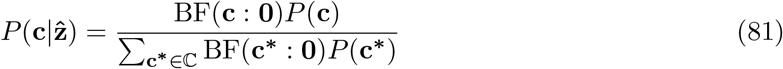

#### 5.3.5 The *ρ*-level credible set

In practice, due to the large number of possible configurations, the probability of any given configuration will likely be small. For more meaningful probabilities, we calculate the total probability of the possible non-null configurations from a set of markers.

We define 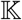 as a set of markers that putatively includes all causal markers. We define 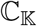 as the set of all causal configurations whose causal markers is a subset of 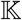, excluding the null set. Thus, the probability that 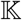 includes all causal markers is the sum of the probabilities of the configurations in 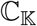.

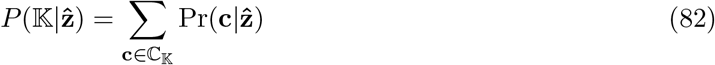

We set this probability as *ρ_c_*, the confidence level of 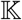. Given a value for *ρ_c_*, commonly 0.95, we seek to find 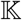 that minimizes the number of causal variants.

#### 5.3.6 The posterior inclusion probability

An alternative way of summarizing the configurations is to calculate a marker’s posterior inclusion probability (PIP), also known as the posterior probability of association. We define the PIP as the probability that a single given marker is causal, marginalized over all other markers. We calculate this probability by summing over all configurations containing the marker.

### 5.4 Computational optimization and implementation

#### 5.4.1 Shotgun stochastic search across configurations

The computation of the probability of a given configuration requires knowledge of the Bayes Factor for every possible configuration. As there are 2^*m*^ possible configurations, traversing this whole space is intractable. To reduce the number of configurations to evaluate, we use the heuristic that configurations with significant probabilities tend to be similar to each other.

We use a shotgun stochastic search procedure based on that of FINEMAP to find all configurations with a signifcant probability [15]. Given a selected configuration **c**, we define 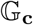, the neighborhood of **c**, as follows:

- All configurations resulting from setting a causal marker in **c** to non-causal
- All configurations resulting from setting a non-causal marker in **c** to causal
- All configurations resulting from swapping the casual statuses of two markers in **c**

For each iteration of the algorithm, the next configuration is drawn randomly from 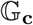, weighted by the posterior probability of each candidate. Upon termination, we assume that all configurations with nonzero probability have been uncovered.

#### 5.4.2 Calculation of the *ρ*-level credible set

Given the large number of configurations evaluated, it is impractical to calculate the best possible credible set satisfying *ρ_c_*. Instead, we use a greedy approximation algorithm utilized in CAVIAR and CAVIARBF [13, 14]. At each step, before *ρ_c_* is reached, the algorithm adds the marker that increases the confidence the most.

#### 5.4.3 Bayes factor evaluation with matrix reduction

Direct calculation of the Bayes Factor for a configuration requires the manipulation of *m × m* matrices, resulting in an *O*(*m*^3^) runtime per configuration. We now show that it is sufficient to evaluate the Bayes Factor using only the elements corresponding to causal SNPs, an optimization introduced by FINEMAP [15]. This reduces complexity from *O*(*m*^3^) to *O*(*k*^3^), where *k* is the number of causal variants.

We expand the MVN probability density functions in equation 80 and use the binomial inverse theorem:

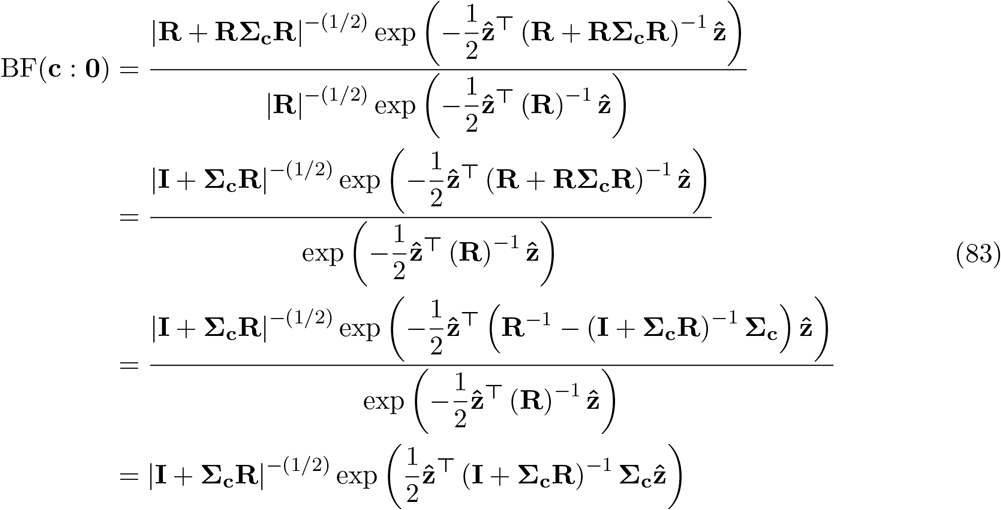

We permute *s* to separate causal and non-causal SNPs:

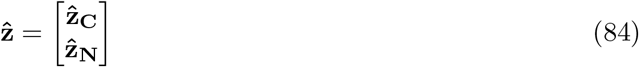

We likewise permute the rows and columns of **R** and **Σ_c_** such that:

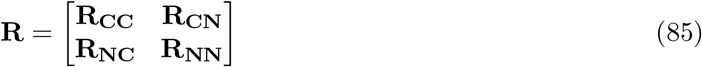

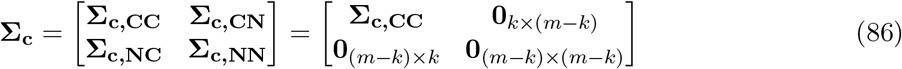

Note that **Σ_c_** can be nonzero only among causal markers since **c** is 0 for non-causal markers. Furthermore:

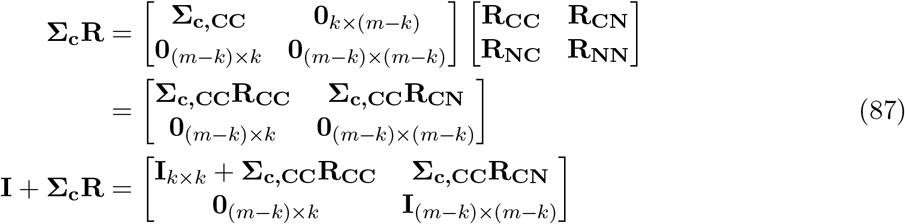

Blockwise inversion yields:

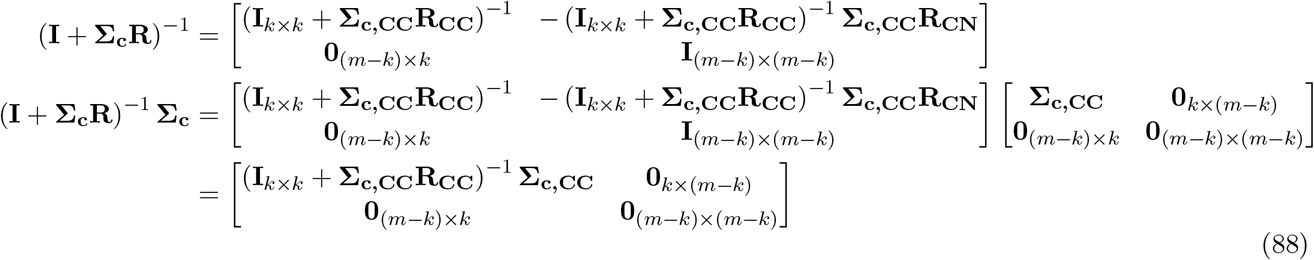

We can simplify this equation since **Σ_c,CC_** is of full rank and is thus invertible:

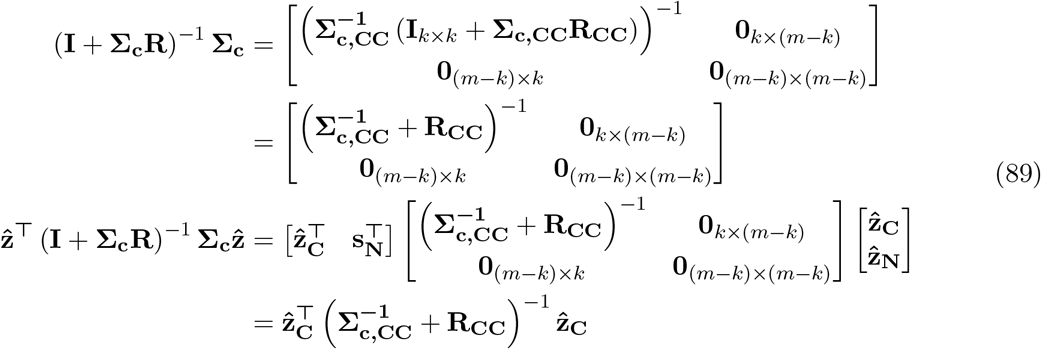

We can also simplify the determinant in Equation 83:

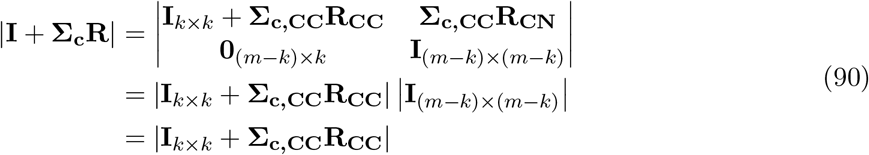

We have thus shown that evaluating the Bayes Factor with only putative causal markers is mathematically equivalent to evaluating with all markers. Thus:

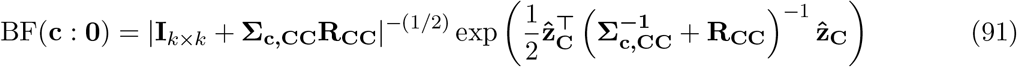

### 5.5 The hyperparameters in terms of heritability

The model takes a number or hyperparameters specifying the variances and covariances of the association statistics. We reparameterize the hyperparameters in terms of the QTL heritablity of the locus 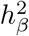 and the AS heritability of the locus 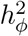.

First, we look at 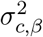 and 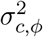, given *k* expected causal variants. Let 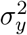 be the overall variance of the QTL phenotype across the individuals. Since the heritability *h_β_* is the proportion of the variance attributed to the causal variance, the average variance of a causal marker’s QTL effect size is given by:

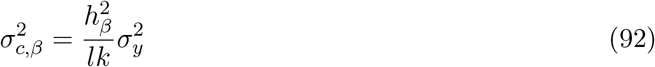

where *l* is the average LD score between the given marker and the other causal markers. Similarly, the variance of the AS effect size is given by:

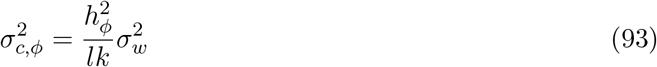

However, in the case of the AS phenotype, where the quality of data varies considerably among individuals, we must take into account the variance introduced by sampling. As we recall, the variance of the observed phenotype for a given individual *j* under a beta-binomial model is:

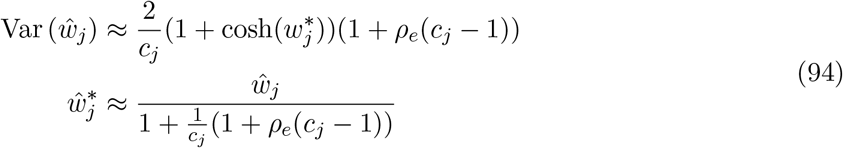

We now derive an estimator 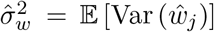 for the total expected variance of the observed phenotype, across individuals. As an approximation, we substitute the individual read coverage for the expected read coverage 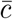, assumng that 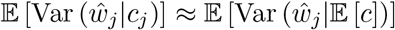. Thus:

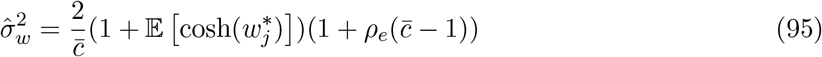

Since we model *ŵ_j_* as normally distributed, 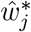 is approximately normally distributed. We now find 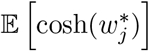. Given a normally distributed zero-mean variable 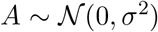, the even-numbered moments are given by:

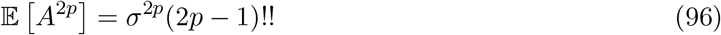

Taking the Taylor expansion of cosh and using linearity of expectation:

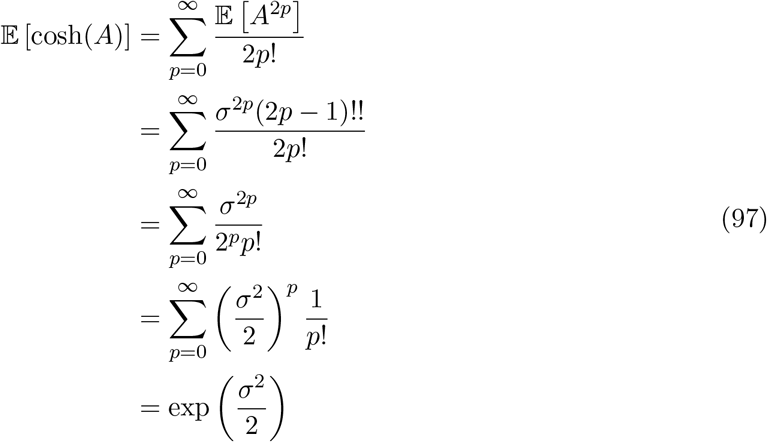

Substituting this result back into the formula for 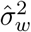:

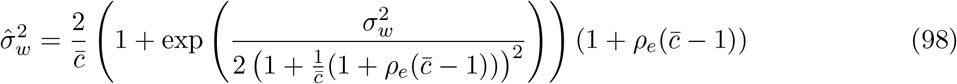

Thus, the variance of the calculated AS effect size is given by:

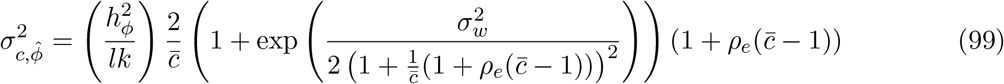

We also define the observed AS heritability 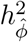 such that

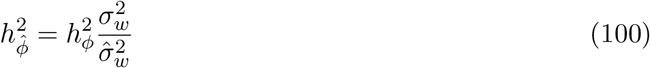

We now derive an expression for *r_βϕ_*, the overall correlation between the QTL and AS statistics for a casual variant. We first find an expression, given causal variant *i* for the variance of 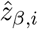. Since *i* is causal, the variance of 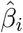 is a combination of the variance of a causal variant and the average phenotypic variation across individuals:

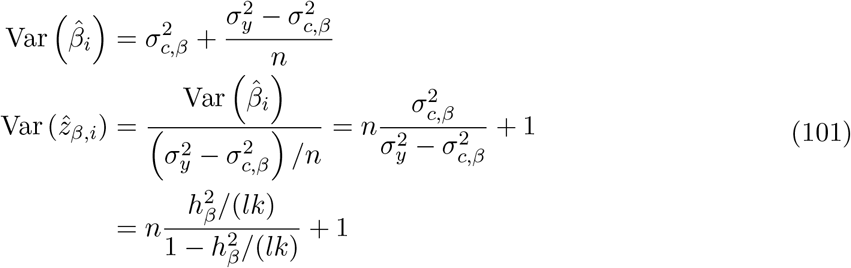

We model the QTL statistic of a causal variant as a zero-mean normal distribution:

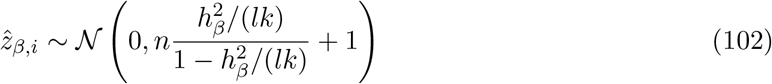

We model the AS statistic in a similar manner:

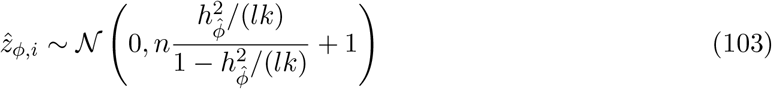

We model the noise as independently distributed between the two statistics, but the causal variance as correlated with coefficient *r_c,βϕ_*. Thus:

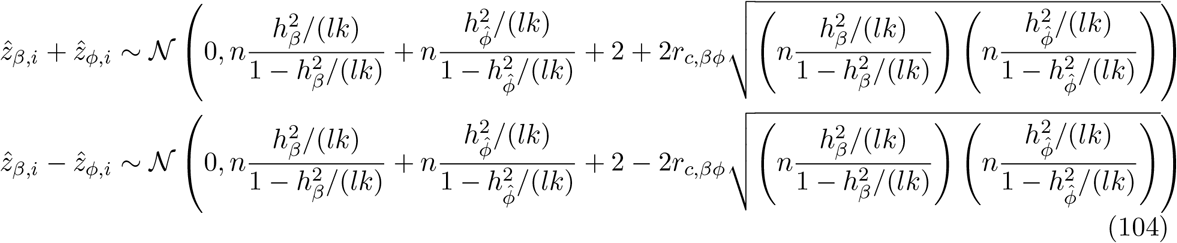

We now find the covariance between 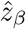 and 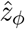. Since the distributions are zero-mean, the covariance is just 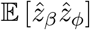. Expanding out this product:

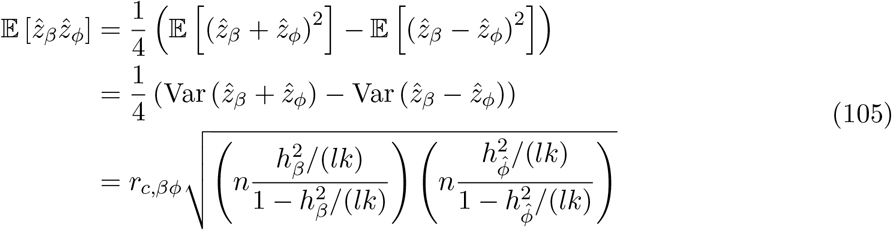

The correlation is thus:

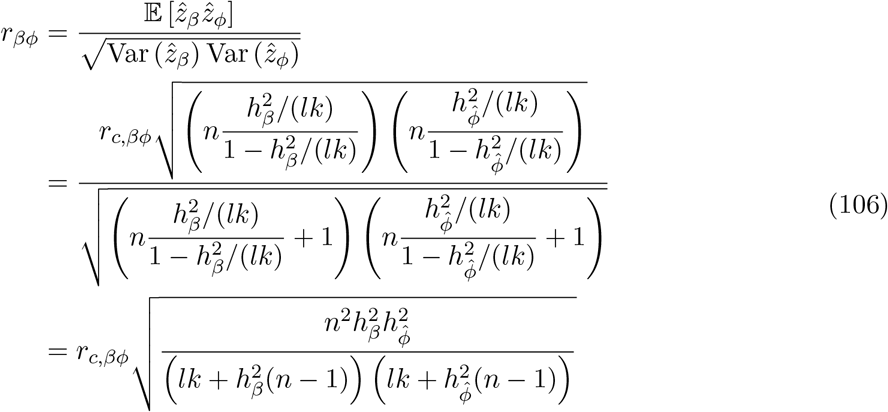

---

**Figure S1:**
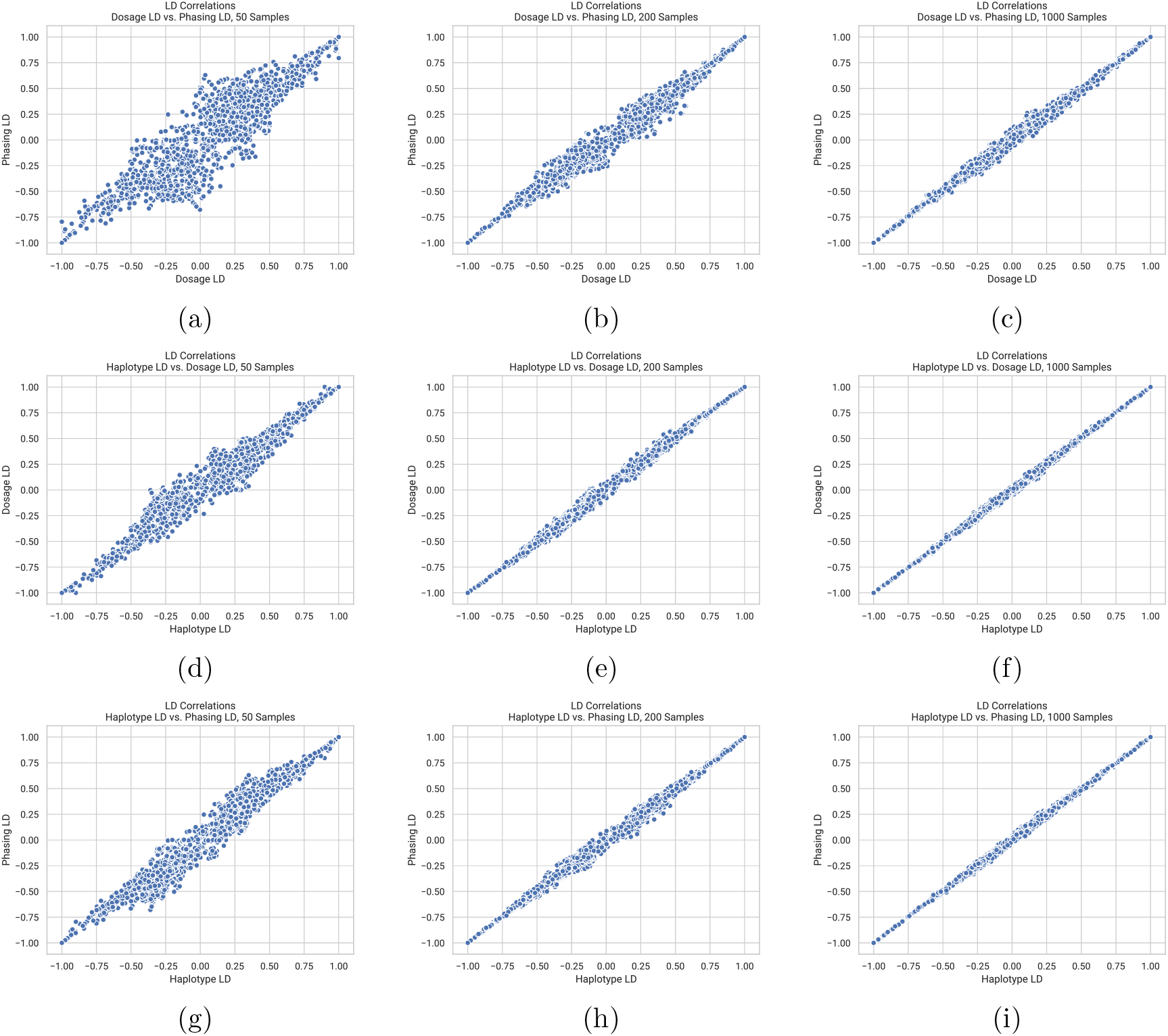
Comparison of estimators for linkage disequilibrium (LD) at various sample sizes. Estimators used were the dosage estimator, the phasing estimator, and the haplotype-level estimator. At each sample size, 100 markers from 1000-Genomes data were used to calculate LD using all three estimators. Plots of LD correlations are shown for each pair of estimators. (a-c) Dosage LD vs. phasing LD at 50, 200 and 1000 samples, respectively. (d-f) Haplotype-Specific LD vs. dosage LD at 50, 200 and 1000 samples, respectively. (g-i) Haplotype-Specific LD vs. phasing LD at 50, 200 and 1000 samples, respectively.

**Figure S2:**
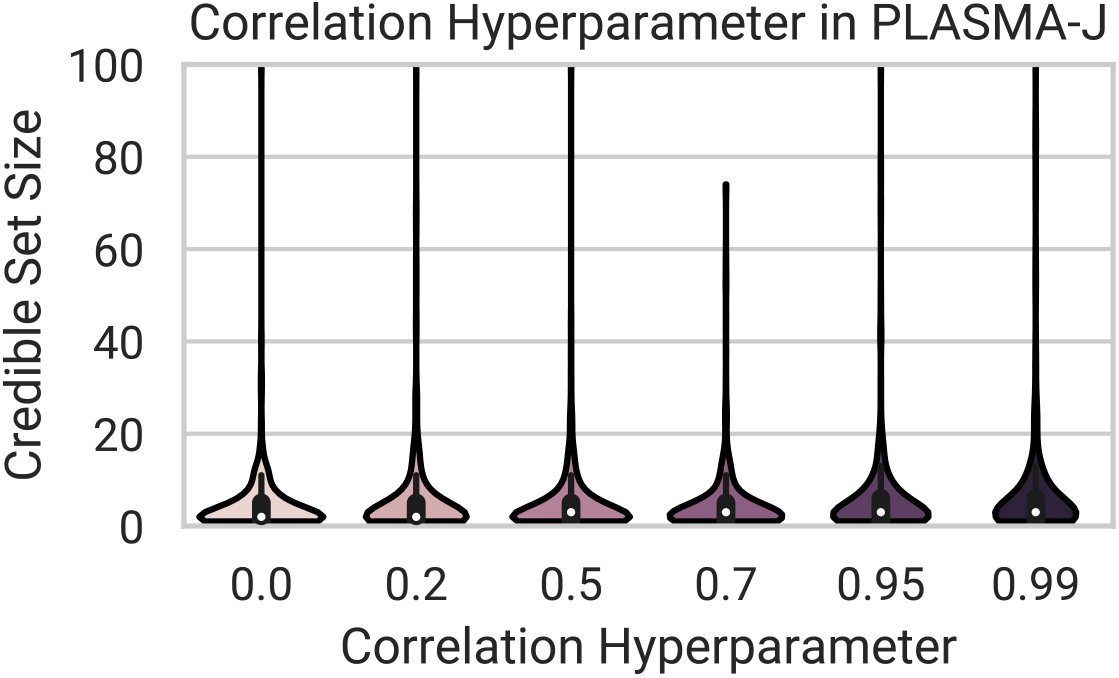
Effect of the jointness hyperparameter in PLASMA on fine-mapping. Fine-mapping of simulated loci was conducted with PLASMA-J under various values of the jointness parameter. Each value of the jointness parameter was evaluated using 500 simulated loci with one causal variant per locus. Plotted are the distributions of the 95% confidence credible sets.

**Figure S3:**
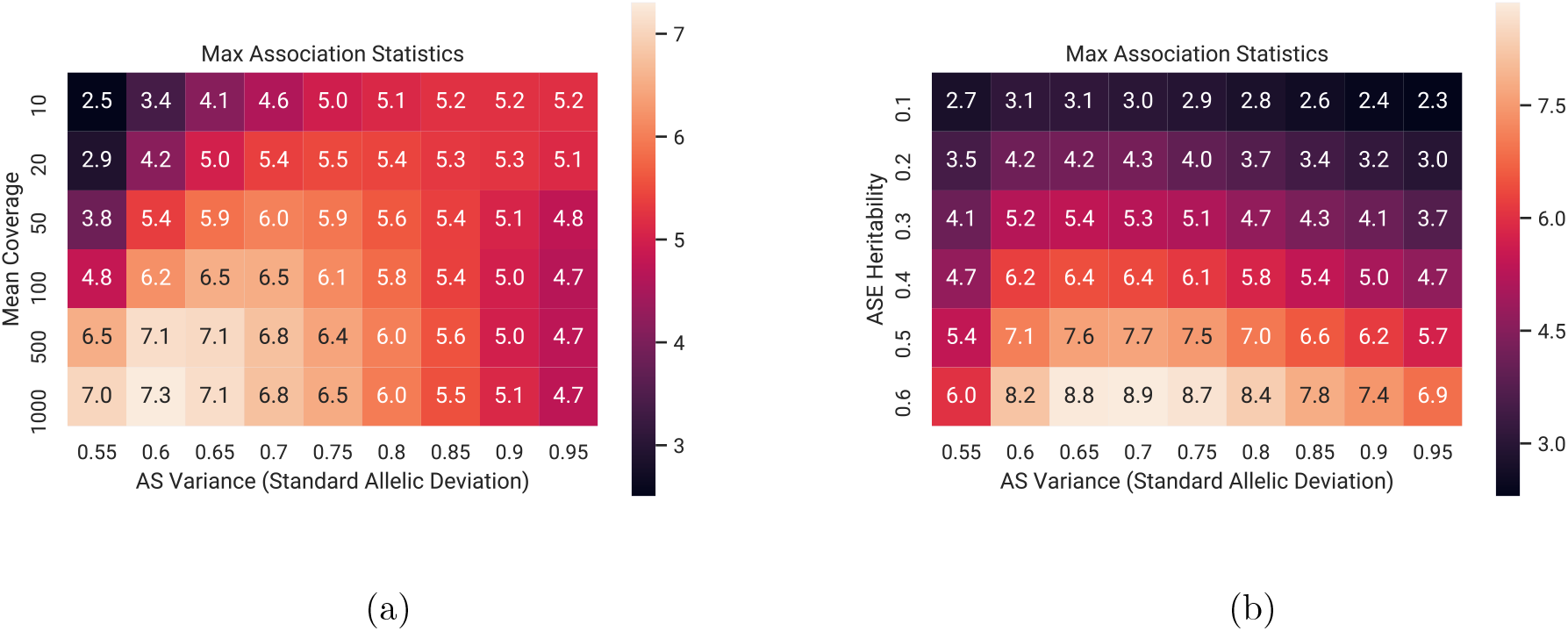
Mean top AS association statistics (*z_ϕ_*) under various conditions of mean read coverage, heritability, and standard allelic deviation (a metric of the variance of the AS phenotype). Standard allelic deviation is defined as the allelic fraction *f* such that the range between 1 – *f* and *f* captures all loci with an AS phenotype within one standard deviation of the mean (balanced expression). Each square is the mean statistic over 500 simulated loci of 100 markers, with one causal variant. (a) Mean top *z_ϕ_* as a function of standard allelic deviation and mean read coverage, with AS heritability set to 0.5. (b) Mean top *z_ϕ_* as a function of standard allelic deviation and AS heritability, with mean read coverage set to 100 reads.

**Figure S4:**
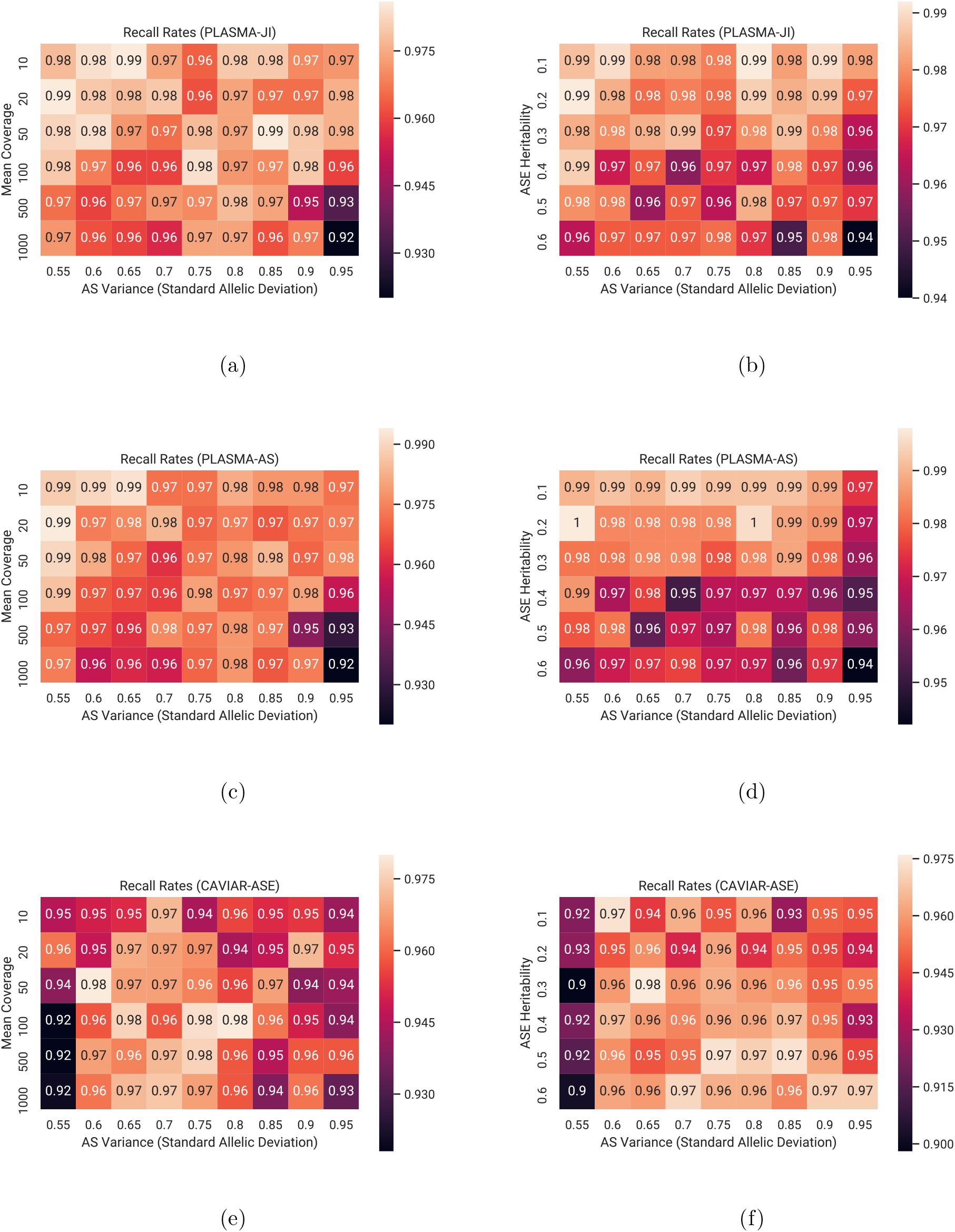
Recall rates for 95% credible sets under various conditions of mean read coverage, heritability, and standard allelic deviation (intensity of AS variance). Recall rate is defined as the expected proportion of causal markers included in the credible set. Each square is the mean statistic over 500 simulated loci of 100 markers, with one causal variant. (a, c, e) Recall rates as a function of standard allelic deviation and mean read coverage, with AS heritability set to 0.4. (d, f, h) Recall rates as a function of standard allelic deviation and AS heritability, with mean read coverage set to 100 reads.

**Figure S5:**
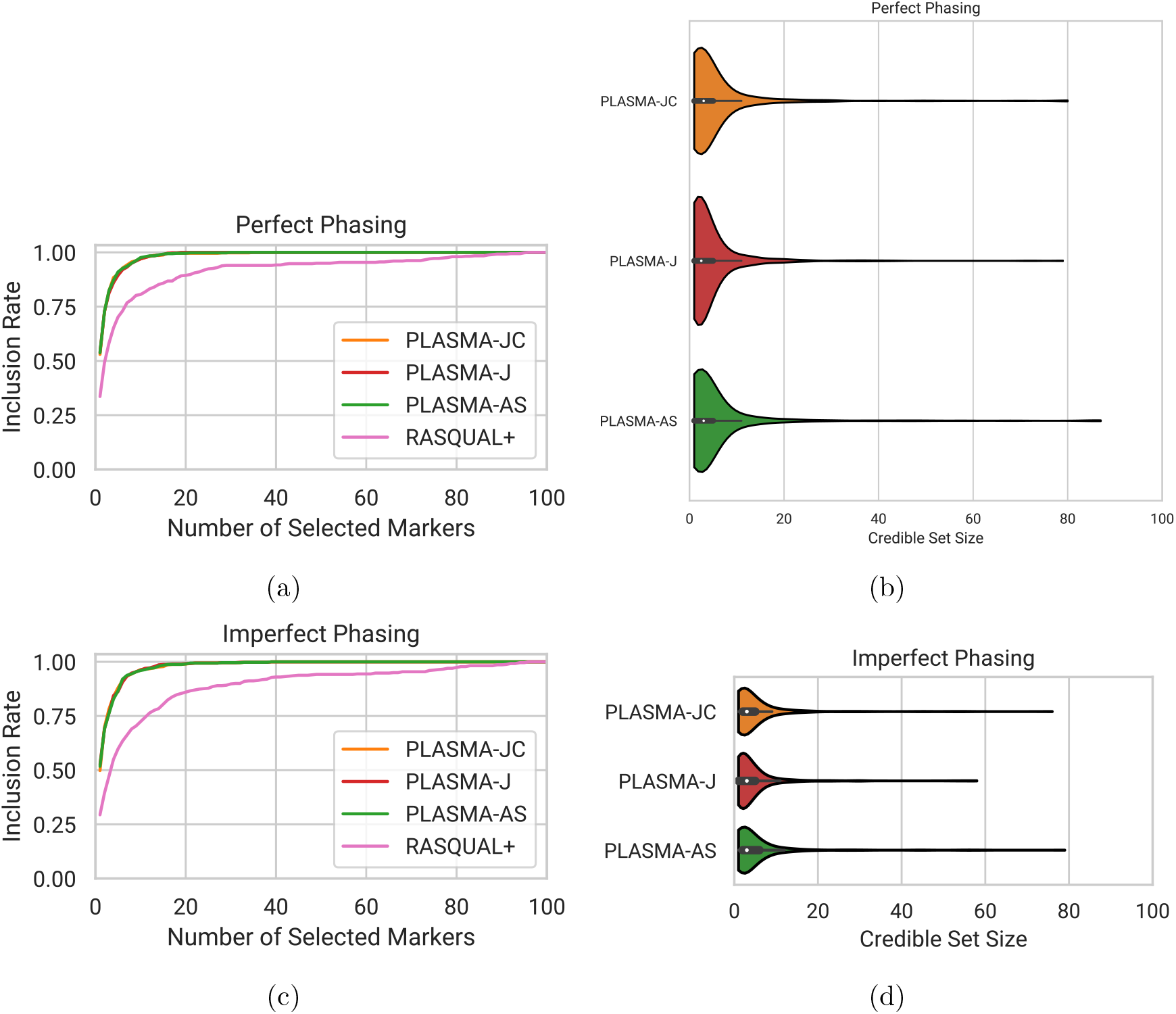
Impact of imperfect phasing on AS-based methods. Each dataset consists of 500 loci with one causal variant per locus. (a, b) Recall and 95% confidence credible sets under perfect phasing. (c, d) Recall and 95% confidence credible sets under imperfect phasing, with a phasing error z-score correction. The switch error and blip error were set to 0.152% and 0.165%, respectively.

**Figure S6:**
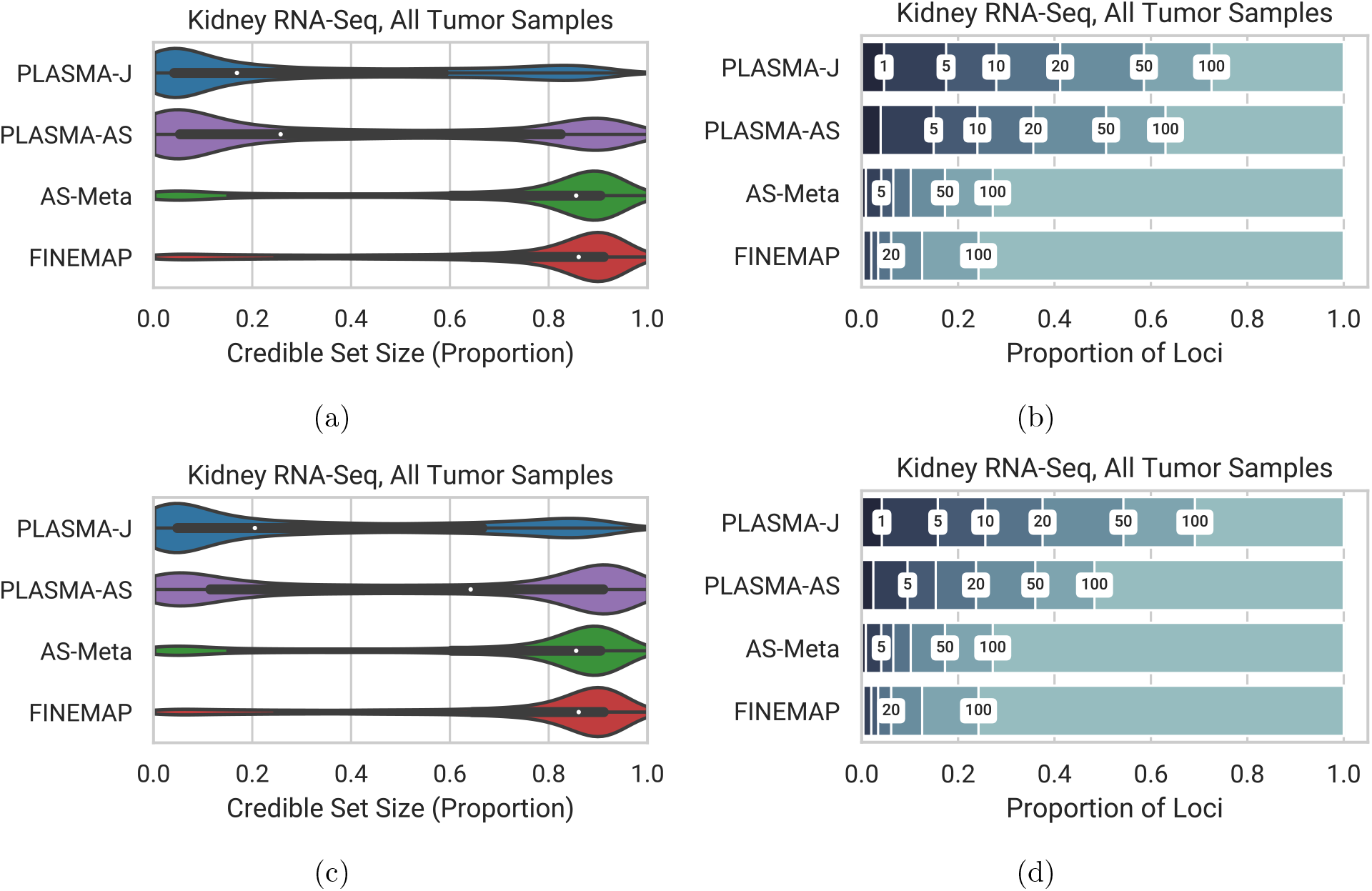
Effect of the AS heritability hyperparameter on 95% credible set sizes with tumor RNA-Seq data. (a) Set size distribution with the heritability hyperparameter at 50% (b) Cumulative set size distribution with the heritability hyperparameter at 50% (c) Set size distribution with the heritability hyperparameter at 5% (d) Cumulative set size distribution with the heritability hyperparameter at 5%

**Figure S7:**
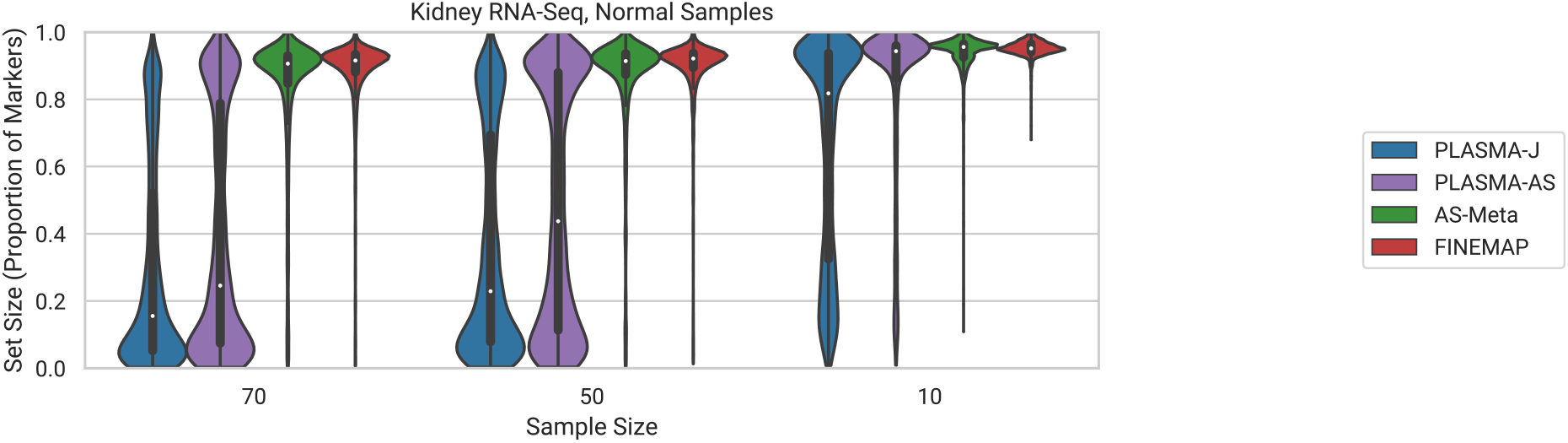
95% credible set size distributions for randomly down-sampled kidney normal data with decreasing sample sizes.

**Figure S8:**
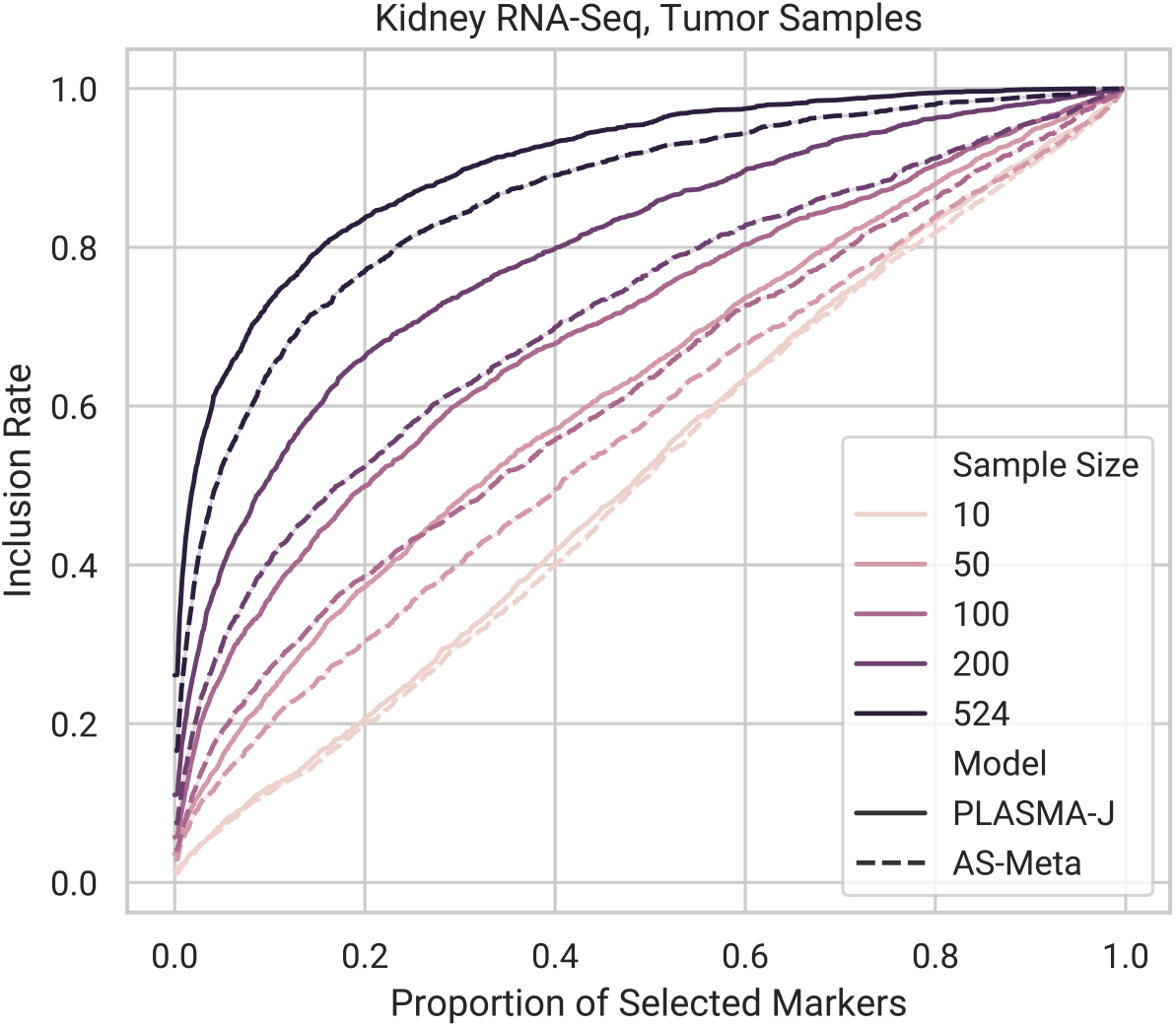
Inclusion curve for fine-mapped kidney tumor RNA-Seq data for PLASMA and AS-Meta across sample sizes. Inclusion was evaluated against a gold-standard set of markers, defined as all markers with at least 0.1 posterior probability under FINEMAP with all samples.

**Figure S9:**
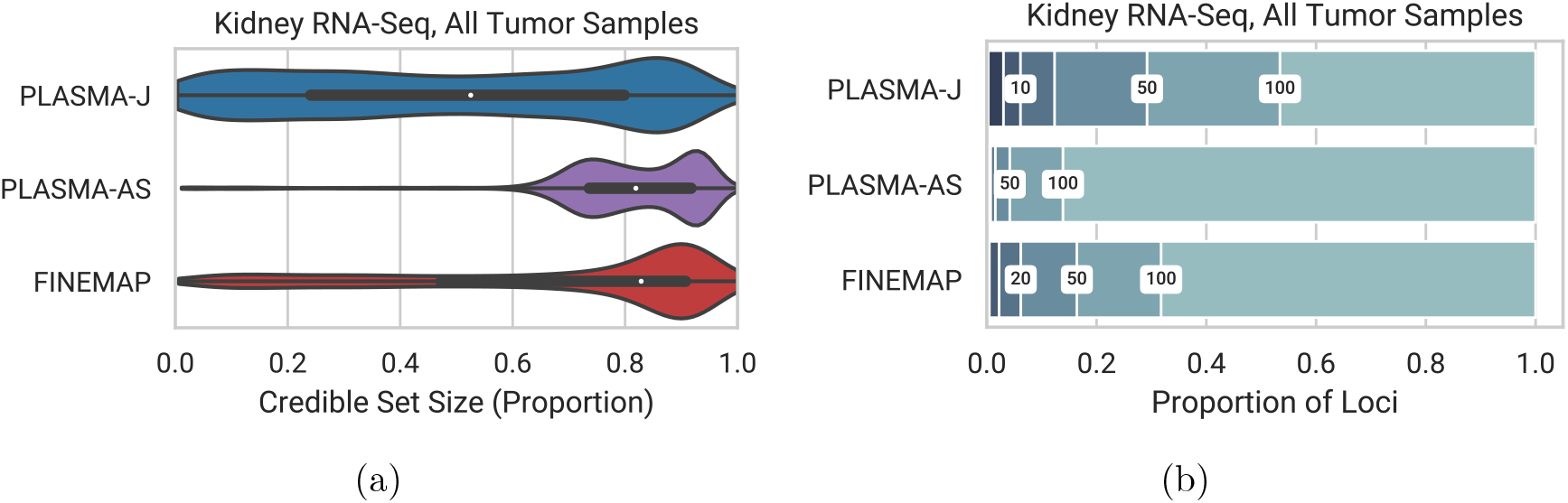
Fine-mapping of kidney RNA-Seq loci under the assumption of up to three causal variants. (Note that FINEMAP credible sets may not be calibrated under multiple causal variants.) (a) 95%confidence credible sets. (b) Proportions of loci with 95% credible set sizes within given thresholds. Thresholds used were 1, 5, 10, 20, 50, and 100.

**Figure S10:**
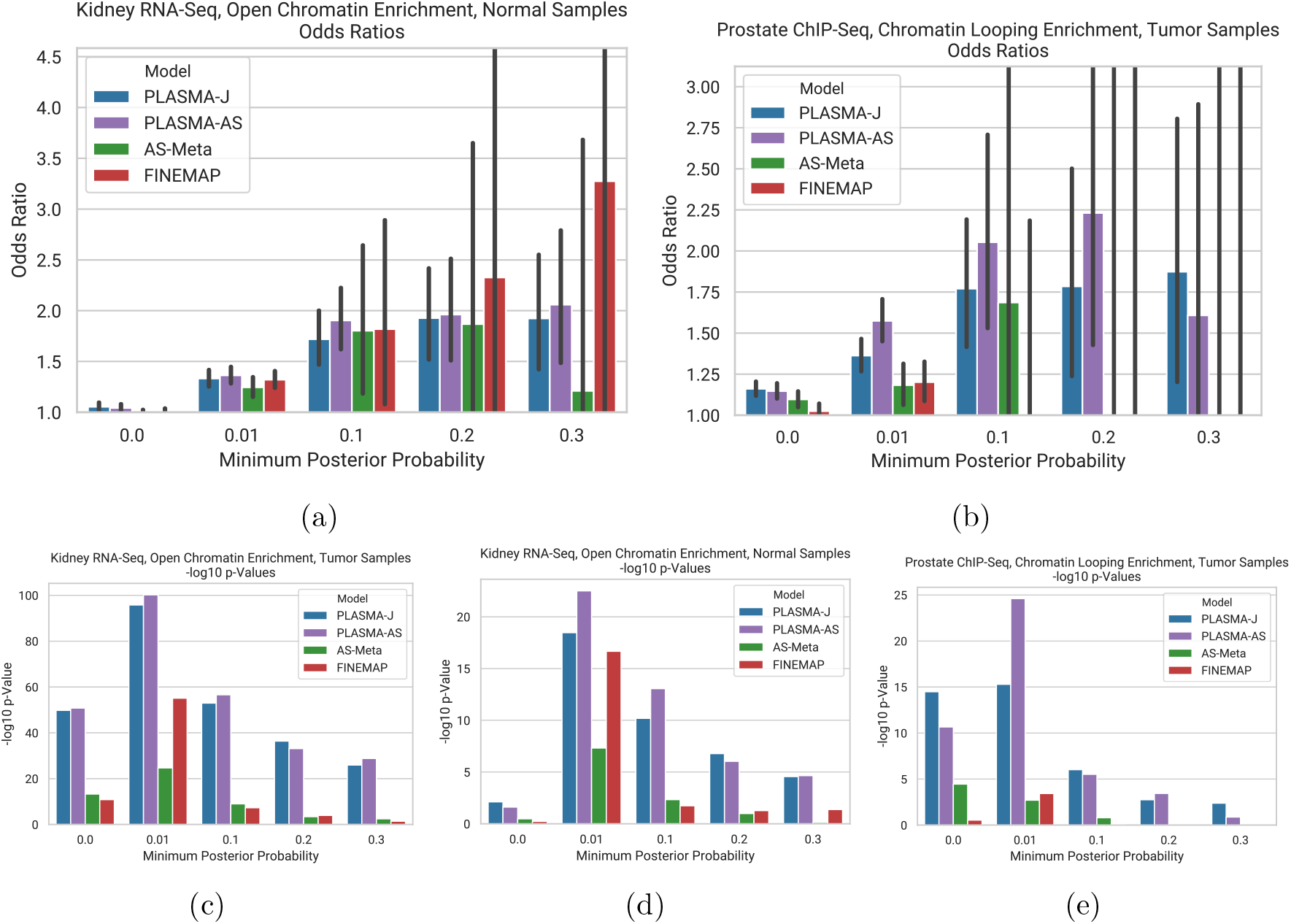
Enrichment odds ratios of PLASMA-J credible sets with functional features. The x-axis represents a minimum posterior probability threshold on the markers within credible sets, with a higher threshold indicating more confident but fewer markers. The markers in these thresholded credible sets were intersected with the functional feature in question. 95 % confidence intervals and p-values were calculated under Fisher’s exact test, with a control of all markers in a locus. (a) Enrichment odds ratios of fine-mapped credible sets with open chromatin for kidney normal RNA-Seq data. (b) Enrichment odds ratios of fine-mapped credible sets with chromatin looping for prostate tumor H3K27ac ChIP-Seq data. (c, d) Enrichment p-values of kidney tumor and normal credible sets, respectively, with open chromatin. (e) Enrichment of fine-mapped credible sets with chromatin looping for prostate tumor H3K27ac ChIP-Seq data.

**Table S1:** Summary of hyperparameter values used in each set of simulation analyses, referenced by figure legend. Parentheses indicate multiple values tested.

| Parameter | Figure 2 | Figures 3a-c | Figures 3d-f |
| --- | --- | --- | --- |
| Number of markers per locus | 100 | 100 | 100 |
| Number of samples | 100 | 100 | 100 |
| Minimum minor allele frequency | 0.01 | 0.01 | 0.01 |
| AS overdispersion ($\rho_e$) | 0.05 | 0.05 | 0.05 |
| QTL Heritability ($h_\beta^2$) | 0.05 | 0.05 | 0.05 |
| AS Heritability ($h_\phi^2$) | 0.4 | 0.4 | (0.1 to 0.6) |
| Manual PLASMA QTL Heritability Setting | N/A | N/A | N/A |
| Manual PLASMA AS Heritability Setting | N/A | N/A | N/A |
| Standard Allelic Deviation ($d$) | (0.6, 0.8) | (0.55 to 0.95) | (0.55 to 0.95) |
| Number of causal variants | 1 | 1 | 1 |
| Mean read coverage | 100 | (10 to 1000) | 100 |
| Min. causal variants in searched configurations | 1 | 1 | 1 |
| Max. causal variants in searched configs | 1 | 1 | 1 |
| Search probability closeness threshold | 0.001 | 0.001 | 0.001 |
| Search iterations convergence threshold | 1000 | 1000 | 1000 |
| Maximum search iterations | 100000 | 100000 | 100000 |
| Jointness parameter ($r_{c,\beta\phi}$) | 0. | 0. | 0. |
| Phasing switch error | 0. | 0. | 0. |
| Phasing blip error | 0. | 0. | 0. |

| Parameter | Figure 4 | Figure S2 | Figure S5 |
| --- | --- | --- | --- |
| Number of markers per locus | 100 | 100 | 100 |
| Number of samples | 100 | 100 | 100 |
| Minimum minor allele frequency | 0.01 | 0.01 | 0.01 |
| AS overdispersion ($\rho_e$) | 0.05 | 0.05 | 0.05 |
| QTL Heritability ($h_\beta^2$) | 0.05 | 0.05 | 0.05 |
| AS Heritability ($h_\phi^2$) | 0.4 | 0.4 | 0.4 |
| Manual PLASMA QTL Heritability Setting | 0.02 | N/A | N/A |
| Manual PLASMA AS Heritability Setting | 0.2 | N/A | N/A |
| Standard Allelic Deviation ($d$) | 0.7 | 0.7 | 0.7 |
| Number of causal variants | 2 | 1 | 1 |
| Mean read coverage | 100 | 100 | 100 |
| Min. causal variants in searched configurations | 1 | 1 | 1 |
| Max. causal variants in searched configs | 3 | 1 | 1 |
| Search probability closeness threshold | 0.001 | 0.001 | 0.001 |
| Search iterations convergence threshold | 1000 | 1000 | 1000 |
| Maximum search iterations | 100000 | 100000 | 100000 |
| Jointness parameter ($r_{c,\beta\phi}$) | 0. | (0. to 0.99) | 0. |
| Phasing switch error | 0. | 0. | 0.00152 |
| Phasing blip error | 0. | 0. | 0.00165 |

